# *In silico* design and validation of high-affinity RNA aptamers for SARS-CoV-2 comparable to neutralizing antibodies

**DOI:** 10.1101/2025.06.04.657802

**Authors:** Yanqing Yang, Lulu Qiao, Yangwei Jiang, Zhiye Wang, Dong Zhang, Damiano Buratto, Liquan Huang, Ruhong Zhou

## Abstract

Nucleic acid aptamers hold promise for clinical applications, yet understanding their molecular binding mechanisms to target proteins and efficiently optimizing their binding affinities remain challenging. Here, we present CAAMO (Computer-Aided Aptamer Modeling and Optimization), which integrates *in silico* aptamer design with experimental validation to accelerate the development of aptamer-based RNA therapeutics. Starting from the sequence information of a reported RNA aptamer, Ta, for the SARS-CoV-2 spike protein, our CAAMO method first determines its binding mode with the spike protein’s receptor binding domain (RBD) through a multi-strategy computational approach. We then optimize its binding affinity via structure-based rational design. Among the six designed candidates, five were experimentally verified and exhibited enhanced binding affinities compared to the original Ta sequence. Furthermore, we directly compared the binding properties of the RNA aptamers to neutralizing antibodies, and found that the designed aptamer Ta^G34C^ demonstrated a comparable binding affinity to the RBD compared to the representative neutralizing antibodies analyzed in this study. This highlights its potential as an alternative to existing COVID-19 antibodies. Our work provides a robust approach for the efficient design of a relatively large number of high-affinity aptamers with complicated topologies. This approach paves the way for the development of aptamer-based RNA diagnostics and therapeutics.

## 1. Introduction

Aptamers are short, single-stranded RNA or DNA oligonucleotides that bind tightly and specifically to target molecules [1,2], earning the designation of “chemical antibodies” [3,4]. They combine the advantageous properties of small organic compounds and antibodies, exhibiting strong affinity and specificity comparable to monoclonal antibodies, while remaining mostly non-immunogenic and highly capable of tissue penetration similar to those of small molecules [5]. These versatile attributes have led to their widespread applications in biomedical fields, including drug discoveries [6], biomarker identifications [7], therapeutics [8], diagnostics [9], and biosensors [10]. The U.S. Food and Drug Administration’s (FDA) approval of Macugen (pegaptanib sodium, Pfizer / Eyetech) in 2004, which targets vascular endothelial growth factor (VEGF) to treat wet age-related macular degeneration, was a landmark achievement for aptamer-based drugs [11]. This success has driven further exploration of aptamer-based therapies. To date, a variety of aptamers have been proposed to neutralize a range of disease-related proteins, such as human epidermal growth factor receptor 2 (HER2) [12], epidermal growth factor receptor (EGFR) [13], and prostate-specific membrane antigen (PSMA) [14]. Recently, the significant contributions of mRNA vaccines [15], aptamers [16] and other RNA biotherapeutics [17,18] have highlighted the potential of RNA-based interventions during the COVID-19 pandemic. This global health crisis has underscored the importance of RNA therapeutics, including medical aptamers.

The systematic evolution of ligands by exponential enrichment (SELEX) technique, the most commonly used method for aptamer screening, still faces several challenges [19]. Firstly, due to its random selection, the candidate aptamers are identified regardless of any atomic mechanisms underlying the interactions between the nucleic acids and target proteins; and secondly, because of limited sizes of oligonucleotide libraries, the identified candidates may not be the best aptamers, which, instead, can be used as lead sequences for further optimization [20]. Therefore, developing *in silico* methods that can complement SELEX screening and provide atomic structure-based rational aptamer design is highly desirable. For instance, we successfully developed two optimized RNA aptamers targeting epithelial cellular adhesion molecule (EpCAM) using a structure-based *in silico* method. We further confirmed their enhanced affinities compared to a previously patented nanomolar aptamer [21]. These two EpCAM-targeted aptamers were relatively small (19 nucleotides) and structurally simple, with a 4-bp stem and a 10-nucleotide long hairpin loop. Their simplicity allowed for precise determination of the aptamer-protein complex structure and enabled structure-based design through conventional computational modelling and simulations. In comparison, RNA aptamers screened through SELEX technique are typically much larger, often comprising tens of nucleotides and featuring complex topologies, including tertiary structural motifs such as internal loops and G-quadruplex. These features not only contribute to increased difficulties in accurate structure modelling, but also increase the possibilities of conformational changes upon binding to the target molecules (i.e. folding upon binding) [22]. Furthermore, the limited number of experimentally determined structures for aptamer-protein complexes in the Protein Data Bank (PDB) present further challenges for an accurate structure modelling. These limitations highlight the need for more advanced computational frameworks to accurately predict the binding modes of RNA aptamers of varying sizes and topologies, thus enabling efficient *in silico* aptamer engineering.

A direct comparison between aptamers (also known as “chemical antibodies”) and conventional biological antibodies in terms of binding mechanisms and affinities is of great interest. However, such comparative studies are currently limited. A prerequisite for such comparison is the availability of a target protein of functional importance for which both effective aptamers and antibodies exist. The receptor binding domain (RBD) of the spike (S) protein expressed by the SARS-CoV-2 is one such target (see Fig. 1A). It binds to human angiotensin-converting enzyme 2 (ACE2) with high affinity to facilitate viral entry into host cells [23,24]. Targeting RBD to block the interactions between SARS-CoV-2 spike protein and human ACE2 emerged as a promising therapeutic strategy [25] to fight COVID-19 pandemic. For example, RBD-targeted antibodies have shown substantial therapeutic potential due to their potent neutralizing effects [26,27]. Alternatively, a recent study reported an RNA aptamer generated through SELEX, termed RBD-PB6-Ta (hereafter referred to as “Ta”), that binds to the RBD of SARS-CoV-2 spike protein with high affinity and efficiently blocks viral infection at low concentrations (see Fig. 1B) [16]. This aptamer was originally identified through multiple rounds of positive and negative SELEX and subsequently validated using surface plasmon resonance (SPR) and biolayer interferometry (BLI), which confirmed its high affinity and high specificity toward the RBD. Therefore, Ta provides a well-characterized and biologically relevant starting point for structure-based optimization. Based on these observations, the RBD of SARS-CoV-2 spike protein represents an ideal target to directly compare the efficacy of aptamers and antibodies. Such comparisons could enhance our understanding of the aptamer binding mechanism, guiding the rational design of aptamers that can possibly be comparable with or even superior to neutralizing antibodies, thereby complementing the existing means of treatment (see Fig. 1C).

**Fig. 1.**
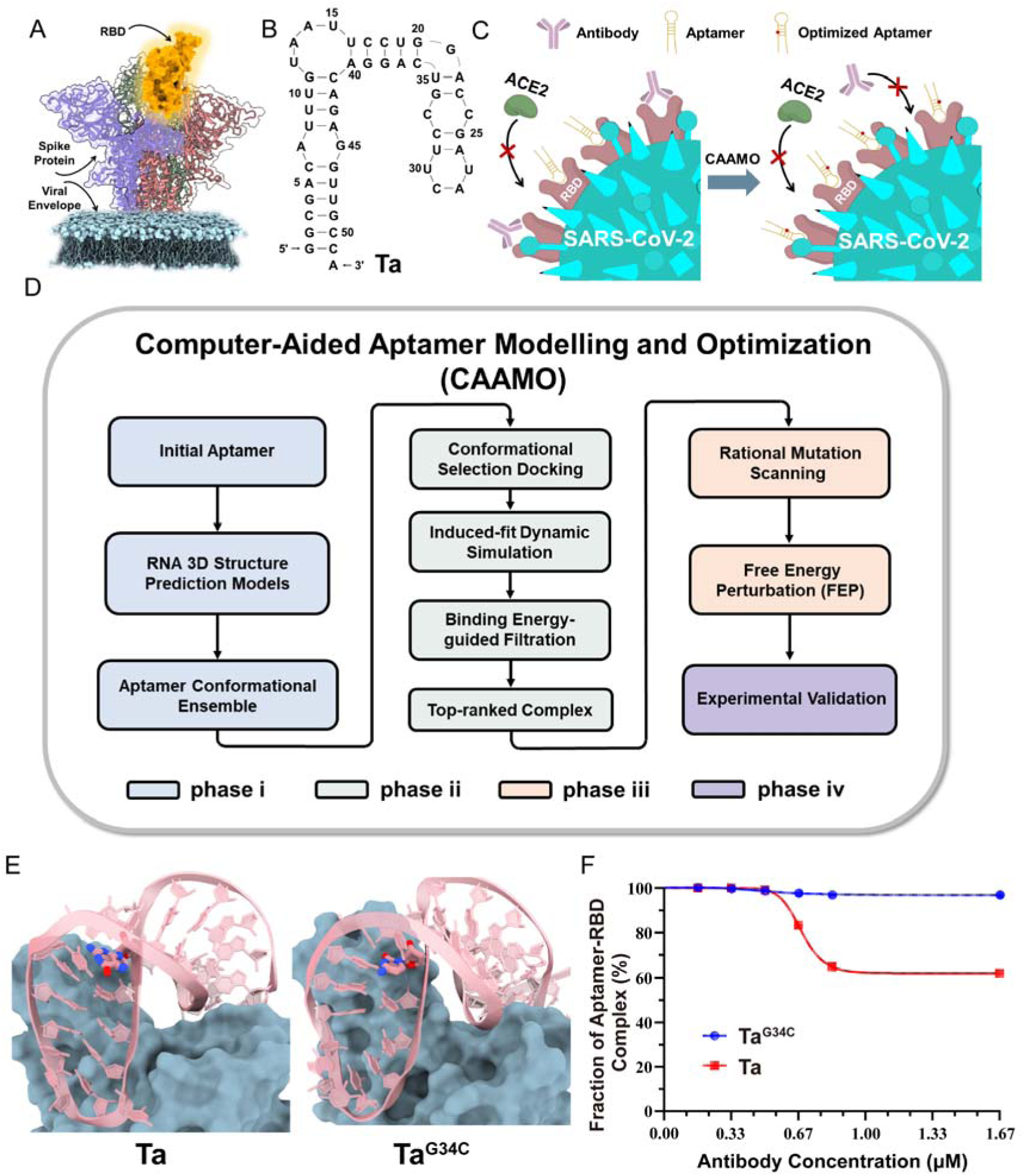
The workflow of the CAAMO framework for designing high-affinity RNA aptamers and its application in the development of novel RNA aptamers targeting the RBD of the SARS-CoV-2 spike protein. (A) Schematic diagram of the spike protein (trimer) of SARS-CoV-2 virus with one of the three RBD regions highlighted in gold. (B) The sequence and Mfold-predicted secondary structure of a SELEX-derived RNA aptamer termed Ta targeting the RBD of SARS-CoV-2 spike protein, which was the initial input information of the CAAMO framework. (C) Schematic diagrams showing that high-affinity RNA aptamers optimized through the CAAMO framework can complement existing antibody-neutralizing treatments for COVID-19. Antibodies (purple) and RNA aptamers (yellow) that bind to the RBDs of the SARS-CoV-2 spike protein can neutralize viral infection by blocking its interaction with the human ACE2. The designed aptamers (marked by red dots) with binding affinities comparable to antibodies can further strengthen the neutralizing treatment when antibody escape occurs. (D) Illustrative workflow of the CAAMO framework by integrating computational techniques with experimental validation. (E) A representative output after CAAMO framework optimization (the aptamer Ta^G34C^) and its comparison with the original Ta sequence. RNA aptamer is colored by pink and RBD is shown in dark blue surface. The mutated nucleotides (G34 in wild type Ta and C34 in optimized Ta^G34C^) are highlighted in pink sticks, atoms oxygen and nitrogen are colored by red and blue, respectively. (F) Competitive binding experiments to compare the binding capabilities of RNA aptamers Ta or Ta^G34C^ and a commercial monoclonal SARS-CoV-2 neutralizing antibody (SinoBiological, Cat: 40592-R001) to the RBD of SARS-CoV-2 spike protein. Gradient amount of the commercial neutralizing antibody (0-1.67 μM) was titrated into the buffer containing 0.5 μM aptamer (Ta or Ta^G34C^) and 40 μM RBD. The intensity of aptamer-RBD bands were quantified with Image J and normalized to that of the mixture without antibody, which was set to 100%. Data were collected from the images of EMSA shown in Fig. 5D.

In this work, we present the CAAMO (Computer-Aided Aptamer Modeling and Optimization) framework, which combines computational techniques and experimental validation to design high-affinity aptamers. Rather than serving as a *de novo* aptamer discovery tool, CAAMO is designed as a post-SELEX optimization platform that rationally improves existing aptamer leads by integrating atomic-level modeling and free-energy–based affinity prediction. Starting with an RNA sequence, we first predicted the most probable binding mode of the RNA aptamer with the target protein using a combination of *in silico* methods, including RNA structure prediction, ensemble docking, molecular dynamics (MD) simulations, steered molecular dynamics (SMD) simulations, and binding free energy calculations. We then performed a comparative analysis between the aptamer Ta and several popular neutralizing antibodies, focusing on their binding sites (using computational methods) and binding capabilities (through both computational and experimental approaches). We identified potential mutation sites for affinity enhancement and developed six novel aptamers through *in silico* mutagenesis study with free energy perturbation (FEP) method. Among these, electrophoretic mobility shift assay (EMSA) experiments confirmed that five had improved binding affinities compared to the original aptamer Ta. Notably, the aptamer Ta^G34C^ exhibited the highest binding affinity to the RBD, outperforming the tested neutralizing antibodies in competitive binding assays. These findings demonstrate the effectiveness of the CAAMO framework in developing high-affinity RNA aptamers targeting the RBD of SARS-CoV-2 spike protein, providing new therapeutic strategies against COVID-19. Moreover, the CAAMO framework can be extended to aptamers that are larger and with more complex topologies.

## 2. Results

**2.1 Overview of the CAAMO framework for high-affinity RNA aptamer design**

For RNA aptamers composed of tens of nucleotides with complicated topological structures, accurately determining their binding modes with target proteins is challenging due to the huge conformational space available, which hinders efforts in structure-based design of high-affinity aptamers. To address these challenges, we propose a promising framework named CAAMO, which integrates computational techniques with experimental validation. This CAAMO framework is designed not only to provide the structural insights into key nucleic acid-protein interactions but also to facilitate the efficient design of aptamers with enhanced affinity. The framework consists of four main phases (see Fig. 1D and Fig. S1): (i) constructing an aptamer conformational ensemble by employing several cutting-edge RNA three-dimensional (3D) structure prediction methods, (ii) identifying an appropriate aptamer binding mode through integrating conformational selection docking, induced-fit dynamic simulation [28], and binding energy-guided filtration, (iii) optimizing the aptamer sequence through *in silico* mutagenesis study using free energy perturbation calculations, and (iv) final experimental validation of the designed aptamer candidates. Further details on the CAAMO framework are provided in the following subsections and in the “Materials and Methods” section.

An RNA aptamer sequence termed “Ta” (containing 52 nucleotides; see Fig. 1B), previously shown to bind the RBD of SARS-CoV-2 spike protein with high affinity [16], was chosen to illustrate the CAAMO framework. The initial input to the CAAMO is the nucleotide sequence of the aptamer Ta, and the output is several *computationally* designed and subsequently *experimentally* validated aptamers with improved binding affinities compared to the initial sequence. A representative output (the aptamer Ta^G34C^), that exhibits a ∼3.3-fold stronger binding affinity than the original aptamer Ta, is shown in Fig. 1E. Notably, the success rate of the *in silico* design via the CAAMO framework is very promising, with five out of the six (∼83%) computationally designed candidate aptamers experimentally confirmed to have improved binding affinities. Furthermore, competitive binding experiments with popular neutralizing antibodies revealed that the designed RNA aptamer Ta^G34C^ has a higher binding affinity to the RBD of SARS-CoV-2 spike protein than the neutralizing antibodies (see Fig. 1F). Thus, the newly identified aptamer Ta^G34C^ shows great potential as a complement to existing antibody-based neutralizing treatments for COVID-19, especially when antibody escape occurs in emerging SARS-CoV-2 variants [29]. Overall, these results indicate that the proposed binding conformation of the aptamer Ta to the RBD serves as a plausible working binding model for structure-guided aptamer optimization, and demonstrate the great potential of our CAAMO framework in aptamer design and optimization.

### 2.2 Determination of the binding model of the aptamer Ta to the RBD

Accurately predicting the binding complex of the RNA aptamer with the target protein is a critical step in structure-based *in silico* aptamer design, especially with aptamers of multiple nucleotides and complex topologies. To address this challenge, we constructed a multi-strategy approach that includes RNA 3D structure prediction, ensemble docking/clustering, and binding capacity assessment to identify the most probable binding conformation of the aptamer Ta with the RBD of SARS-CoV-2 spike protein (see Fig. 2A). The RBD conformation was extracted from the crystal structure of the RBD-ACE2 complex (PDB id: 6LZG) and then refined using MD simulation (see Fig. S2). For the aptamer Ta, we first predicted its secondary structure using Mfold [30], forming a stem-loop structure containing five stems, two internal loops, two bulges and one hairpin loop (see Fig. 1B). Then, we built a 3D conformational ensemble using the state-of-the-art RNA 3D structure prediction models because of the inherent flexibility of unpaired nucleotides in loop regions and potential conformational changes upon binding to the target protein. A total of 25 representative aptamer Ta structures with different degrees of bending (see Fig. S3A) were selected for subsequent molecular docking. It should be noted that, since most popular structure prediction models, such as FARFAR2 [31], IsRNA2 [32] and SimRNA [33], adopt a conformation clustering strategy to obtain the top predictions, each of these 25 predicted structures represents a collection of similar 3D conformations.

**Fig. 2.**
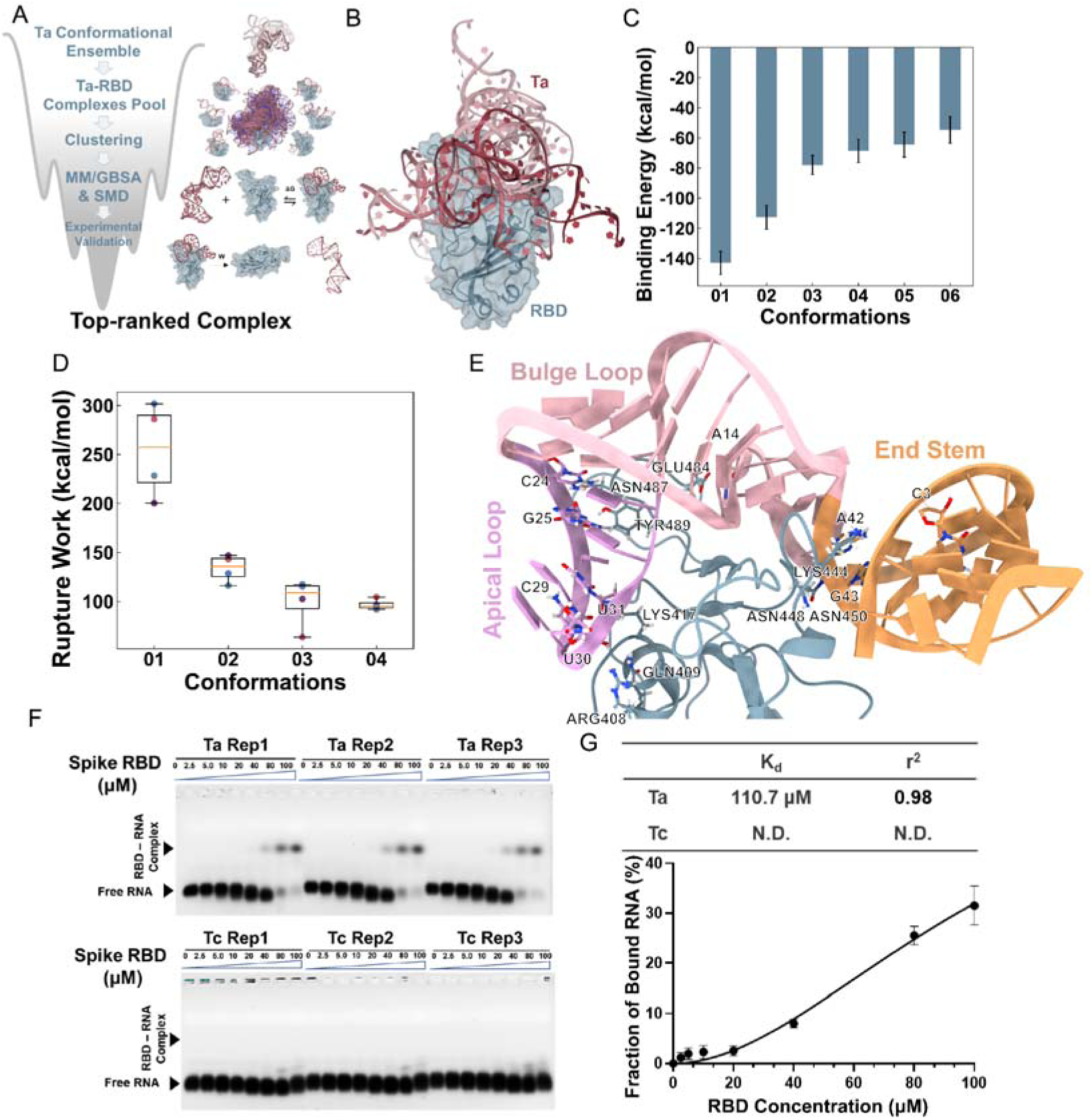
Determination of the binding model of the aptamer Ta and the RBD of the SARS-CoV-2 spike protein *via* a multi-strategy approach. (A) Flowchart illustrating the combination of RNA 3D structure prediction, ensemble docking and clustering, and binding capacity assessment by MM/GBSA and steered MD to determine the most probable binding conformation of the aptamer Ta to the RBD. (B) Six main binding modes, i.e., conformations 01-06, with clearly distinct binding conformations after ensemble docking and clustering for further binding ability assessment. The RBD and the aptamer Ta are shown in dark blue and dark red, respectively. For clarity, the surface of RBD is also displayed. For the aptamer Ta, conformations 01-06 are named in a descending order of their respective cluster sizes and their colors are lightened gradually. (C) The binding energies between the aptamer Ta and RBD estimated by MM/GBSA method for six binding candidates shown in plane (B). Data represent mean±s.d. collected from three independent calculations. (D) The rupture works required to separate the bound aptamer Ta from the RBD for different binding conformation candidates. Data were collected from four independent steered MD simulations. Limited by the available computing resources, only the first four binding conformations were assessed. The orange line is the median, boxes extended from lower to upper quartiles, whiskers showing the range of nonoutlier data. (E) Overview of the most probable binding model (conformation 01) of the aptamer Ta and RBD. The RBD is shown in cyan cartoon representation, the apical loop, bulge part, and end stem of the aptamer Ta are colored plum, pink, and sandy brown, respectively. The key residues of the RBD and nucleotides of the aptamer Ta for binding interactions are shown in sticks, all nitrogen atoms colored by blue, and all oxygen atoms by red. (F) EMSA results of the aptamer Ta (upper panel) and the weakly binding aptamer Tc (negative control, lower panel) bound to the RBD of the SARS-CoV-2 spike protein and (G) the resultant binding curve for the aptamer Ta. The dissociation constant (*K_d_*) was calculated from the EMSA image quantification from three independent experiments (Mean±s.d., n = 3).

From the ensemble docking complex pool (containing 1,000 aptamer-RBD binding poses; see Fig. S3B), we generated six major clusters, each representing distinct binding conformations. For each cluster, we selected the structure with the lowest binding energy, estimated using the molecular mechanics generalized Born surface area (MM/GBSA) approach based on single-frame conformation, as the representative binding conformation (see Fig. 2B and conformations 01-06 in Fig. S3C). MD simulations of these six representative conformations confirmed that they maintained stable binding modes over the course of the long-time simulations (see Fig. S4). We further analyzed the binding abilities between the aptamer Ta and RBD for these six MD refined conformations using MM/GBSA and steered molecular dynamics (SMD) simulations to determine the most likely binding conformation. As shown in Fig. 2C, the MM/GBSA binding energy of conformation 01 (Δ*G*=-142.93±7.65 kcal/mol) is significantly lower than the other five candidates (Δ*G*>-120 kcal/mol). Interestingly, we found that the binding energy of each conformation correlated with the sizes of its corresponding clusters (see Fig. 2C and Fig. S3C), with a larger cluster associated with lower binding energy. Additionally, we computed the rupture works required to separate the bound aptamer from the RBD using SMD simulations. Among four tested cases (conformations 01-04), the conformation 01 rupture work is the largest (254.2±47.9 kcal/mol) and significantly different from that of the other conformations (see Fig. 2D). Despite the limitations of MM/GBSA and SMD methods (more rigorous FEP method will be used later), both approaches consistently showed that conformation 01 had the strongest binding ability between the aptamer Ta and the RBD. Based on these results, we selected conformation 01 as the putative binding model for further structure-based rational design of high-affinity aptamers.

In the putative binding complex, the aptamer Ta adopts a saddle-like shape to bind the RBD (see Fig. 2E). We divided the aptamer into three parts: the apical loop (contains two base pairs; nucleotides C24-C33), the bulge region (nucleotides G11-C23 and G34-C41), and the end stem part (contains one mismatch base pair; nucleotides G1-U10 and A42-A52). The apical loop and end stem parts bind to opposite sides of the RBD, respectively, while the bulge acts as a linker, adjusting the bending angle of the saddle-like shape. The binding between the aptamer Ta and RBD is stabilized by electrostatic, hydrogen bonding, and van der Waals interactions. In details, some important interactions include the electrostatic interactions between the phosphate groups of nucleotides C29 and U30 and the basic amino acid ARG408, and between the phosphate groups of C3, A42 and G43 and the LYS444; the hydrogen bonds between GLN409@Nε2-Hε21…C29@O2’, LYS417@Nζ-Hζ3…U31@O2, U42@O2’-HO2’…ASN448@Oδ1, ASN450@N-H…U42@ O2’, A14@N6-H61…GLY482@O, A14@O2’-HO2’…GLU484@Oε2, ASN487@ Nδ2-Hδ22…C24@O2’ and TYR489@OH-HH…G25@O2’ and the van der Waals packing interactions between nucleotides of the apical loop part and some aromatic residues of the RBD (such as TYR421, TYR453, PHE456, TYR473, PHE486, and PHE489) (see Fig. S5).

These detailed interactions provide important insights for future *in silico* aptamer design. We note that CAAMO is not intended to establish experimentally validated complex structures, but rather to provide preliminary binding models that enable rational affinity maturation of aptamers in scenarios where structural information is limited or unavailable.

In addition to the aptamer Ta, we also analyzed a weaker binding aptamer sequence, RBD-PB6-Tc (Tc), as a negative control [16]. We constructed a binding model for the Tc sequence with the RBD using the same approach as for the aptamer Ta (see Fig. S6). SMD simulations confirmed that the binding strength of Tc to the RBD was significantly weaker than that of the aptamer Ta (see Fig. S6D), which supports the robustness of our approach in generating informative binding models for comparative analysis and affinity optimization of an RNA aptamer with a target protein. Additionally, our EMSA experiments also confirmed that the binding affinity of the Ta-RBD complex is stronger than that of the Tc-RBD complex (see Figs. 2F and 2G), in good agreement with the previous study [16]. The measured dissociation constant for the Ta-RBD complex is *K_d_*=110.7 μM, while the sequence Tc is unable to bind to the RBD under all conditions tested. These results motivate the design of high-affinity RNA aptamers targeting the RBD of the SARS-CoV-2 spike protein to enhance the treatment of COVID-19.

### 2.3 Comparison of binding properties between the aptamer Ta and neutralizing antibodies

In recent years, researchers have developed numerous antibodies to neutralize SARS-CoV-2 infection by blocking the RBD of the spike protein from binding to ACE2 [34]. Their binding properties with the RBD have been extensively studied. Since we determined the most probable binding model of the aptamer Ta to the RBD, comparing the binding properties of the aptamer Ta with those of representative neutralizing antibodies to the RBD is both feasible and meaningful. A preliminary comparison of the respective binding modes of ACE2, the aptamer Ta, and neutralizing antibodies to the RBD (see Fig. S7A) indicates that the aptamer Ta can precisely occupy the binding site of ACE2, similar to many neutralizing antibodies, thereby blocking SARS-CoV-2 invasion. To further explore this, we analyzed the contact ratios of residues on the RBD bound to ACE2 (derived from MD simulations), to the aptamer Ta (derived from MD simulations), or to the neutralizing antibodies (derived from all available experimentally resolved SARS-CoV-2 RBD–antibody complex structures curated in the Coronavirus Antibody Database, CoV-AbDab [35]). CoV-AbDab is a publicly available, curated database that aggregates all published coronavirus-binding antibodies with associated structural information, providing a comprehensive and unbiased structural ensemble for contact frequency analysis. And the results indicate that the contact regions on the RBD bound by the aptamer Ta, ACE2 or the neutralizing antibodies were largely similar (Fig. 3A). Additionally, we analyzed the electrostatic potential distribution on the RBD surface to assess how the highly negatively charged RNA aptamer interacts with the RBD (see Fig. 3A). The positively charged regions on both sides of the RBD promote the aptamer binding. To quantitatively analyze the binding complexes, we defined a key residue as the one with a contact ratio greater than 0.5. We found that ACE2 binds to the RBD using 12 key residues, such as LYS417, LEU455, PHE456, ALA475 (see Fig. 3B) while the neutralizing antibodies engage 19 key residues on the RBD, including all of ACE2’s key residues except for THR500. This higher number of key residues corresponds with the stronger binding ability of neutralizing antibodies compared to ACE2, enhancing their ability to block SARS-CoV-2 invasion. For the aptamer Ta, we identified 27 key residues, most of which overlap with those of ACE2 and neutralizing antibodies. For instance, nine key residues on the RBD are shared by ACE2, neutralizing antibodies, and the aptamer Ta (see Fig. 3B), suggesting that the aptamer Ta may have comparable or even better binding ability to the RBD compared to the neutralizing antibodies.

**Fig. 3.**
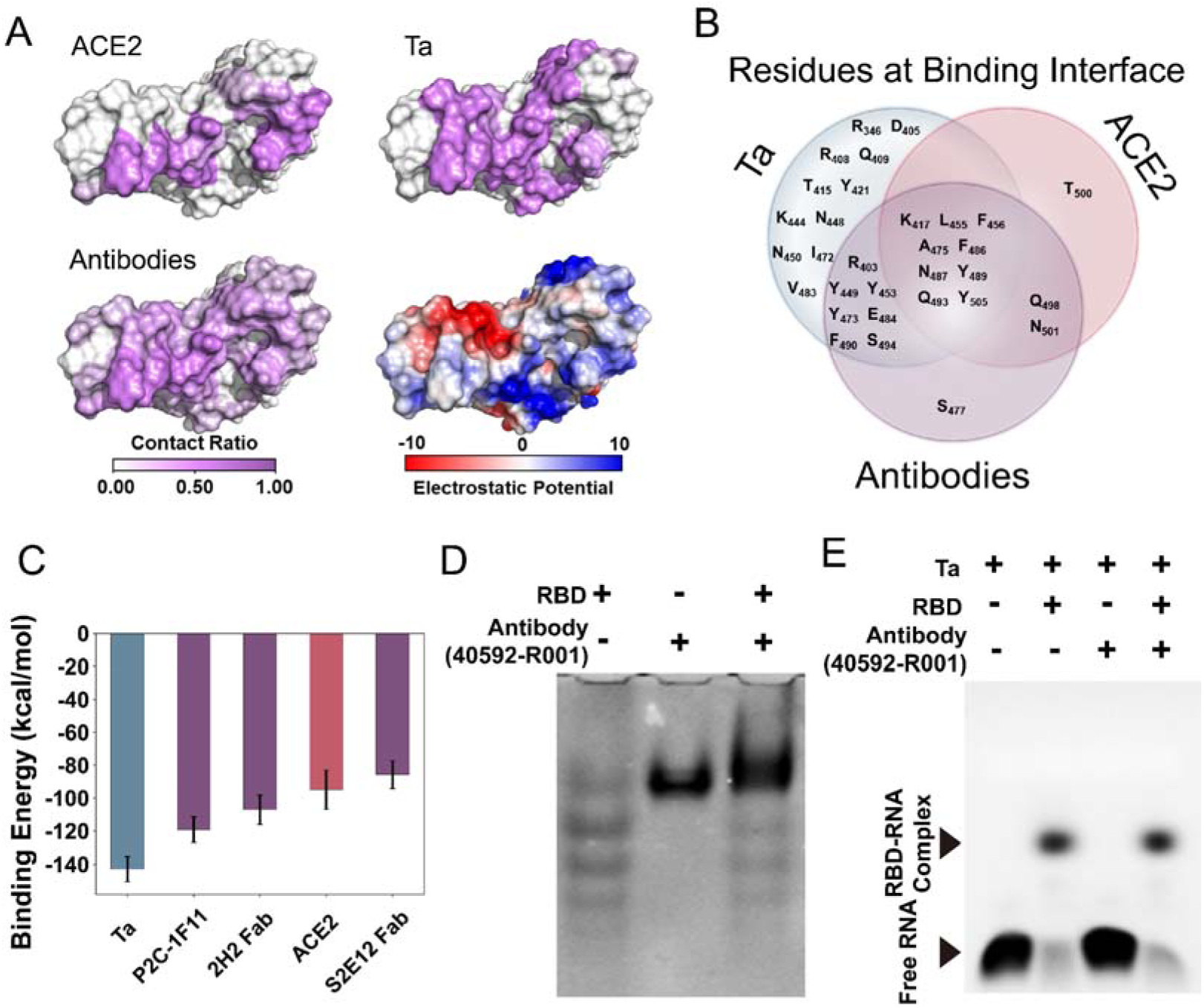
The aptamer Ta exhibits a comparable binding capability to RBD compared to neutralizing antibodies. (A) Contact ratios of residues on the RBD by ACE2 (derived from MD simulations), the aptamer Ta (derived from MD simulations), and neutralizing antibodies (derived from all available SARS-CoV-2 RBD–antibody complex structures curated in CoV-AbDab). For reference, the electrostatic potential distribution on the RBD surface generated by PyMOL (version 2.3.5) program was also shown. (B) Key residues on RBD (with contact ratio larger than 0.5 in plane (A)) contacted by ACE2, the aptamer Ta, and neutralizing antibodies were displayed as a Venn diagram. (C) Binding energies estimated by MM/GBSA calculations for ACE2, the aptamer Ta, and three representative antibodies (P2C-1F11, 2H2, and S2E12) binding to RBD. Data represent mean±s.d. collected from three independent calculations. (D) Binding ability of the commercial antibody (40592-R001) to RBD was assessed by native-PAGE. The reaction mixture (10 μL) contained 8.57 μM RBD protein and 2 μM antibody, incubated individually or combined, followed by Coomassie brilliant blue staining. (E) The RBD binding abilities of the aptamer Ta and commercial antibody 40592-R001 were compared by EMSA competitive binding experiments. The aptamer-RBD complex bands were shown after running on an agarose gel following the incubation of 40 μM RBD protein, 0.5 μM aptamer Ta, and 0.5 μM antibody 40592-R001 (final concentrations in the reaction mixture).

We performed binding energy estimations using MM/GBSA calculation and competitive binding assays with EMSA experiments to further compare the binding abilities of the aptamer Ta and neutralizing antibodies to the RBD. We selected three potent antibodies that can effectively neutralize SARS-CoV-2 infection, including antibody P2C-1F11 [36] (PDB id: 7CDI), 2H2 [37] (PDB id: 7DK4), and S2E12 [34] (PDB id: 7K4N), and calculated their binding energy to the RBD, along with that of ACE2. As shown in Fig. 3C, antibodies P2C-1F11 (-119.24±7.86 kcal/mol) and 2H2 (-107.18±8.89 kcal/mol) have lower binding energies (indicating stronger binding abilities) to RBD than ACE2 (-94.94±11.95 kcal/mol), while antibody S2E12 (-85.77±8.55 kcal/mol) exhibits a similar binding energy to ACE2. These findings confirm the reliability of our binding energy estimations using MM/GBSA calculation. The aptamer Ta, with a binding energy of-142.93±7.65 kcal/mol, has a significant lower binding energy than both ACE2 and the three antibodies, indicating a stronger binding ability of the aptamer Ta to the RBD. We further tested the binding ability of the aptamer Ta with a commercial neutralizing antibody (SinoBiological, Cat: 40592-R001, termed 40592-R001) that targets the RBD of the SARS-CoV-2 spike protein. The antibody 40592-R001 demonstrated high affinity for the RBD in our native gel protein-protein binding assays (Fig. 3D). We then added the antibody 40592-R001 into a mixture of the aptamer Ta and RBD, monitoring the changes in the formation of aptamer-RBD complex via fluorescence intensity in the EMSA experiments. In principle, if the aptamer Ta binds more strongly to RBD than the antibody 40592-R001, the presence of the antibody should not disrupt the formation of the aptamer-RBD complex; otherwise, the amount of the aptamer-RBD complex would be reduced. Notably, the Ta-RBD complex formation remained unchanged after adding the antibody (Fig. 3E), suggesting that the aptamer Ta exhibits binding capability comparable to the tested monoclonal neutralizing antibody. These results also confirmed that the aptamer Ta binds to the same site on the RBD as the 40592-R001 antibody. In conclusion, the aptamer Ta presents a promising alternative, or at least a complementary approach, to conventional antibody-based neutralizing therapies against COVID-19. These findings prompt us to further optimize the aptamer Ta to enhance its binding ability to the RBD of the SARS-CoV-2 spike protein.

### 2.4 Structure-based rational design of the aptamer Ta to improve binding affinity

Since the SELEX screening process explored only a limited sequence space, we anticipated that the SELEX-derived aptamer Ta could be further optimized *via* rational mutation scanning to improve its binding affinity to the RBD of the SARS-CoV-2 spike protein. In principle, nucleotide mutations in the aptamers can affect the binding affinity, which can be characterized by changes in binding free energy (ΔΔ*G*), and we accurately assessed these changes using FEP calculations. A negative change in binding free energy (ΔΔ*G*<0) indicates improved binding affinity, while a positive change (ΔΔ*G*>0) means a decrease in binding affinity. FEP is regarded as one of the most rigorous and reliable methods in estimating binding free energy changes, achieving high accuracy in identifying key residues and their mutational effects for many protein-protein, protein-ligand, and protein-RNA complexes, with results comparable with experiments [21,38]. However, unlike protein and peptide drugs, the structure of RNA molecules is very sensitive to its nucleotide composition and even a single mutation may cause significant changes in its secondary (2D) structure, potentially affecting its binding modes. To address this, we employed secondary structure analysis (SSA) to examine the structural similarity before and after nucleotide mutation. Moreover, we used MFold [30] to predict the 2D structure with the lowest free energy for each mutated sequence. Structural similarity between the wild-type (WT) Ta sequence and its mutants was quantified by base-pair (BP) similarity, defined as the ratio of shared base pairs (*N_shared_*) to the total base pairs in the WT sequence (*N_WT_*). Overall, as summarized in Fig. 4A, we combined rational mutation scanning, SSA, and FEP to design Ta analogues with enhanced binding affinity.

**Fig. 4.**
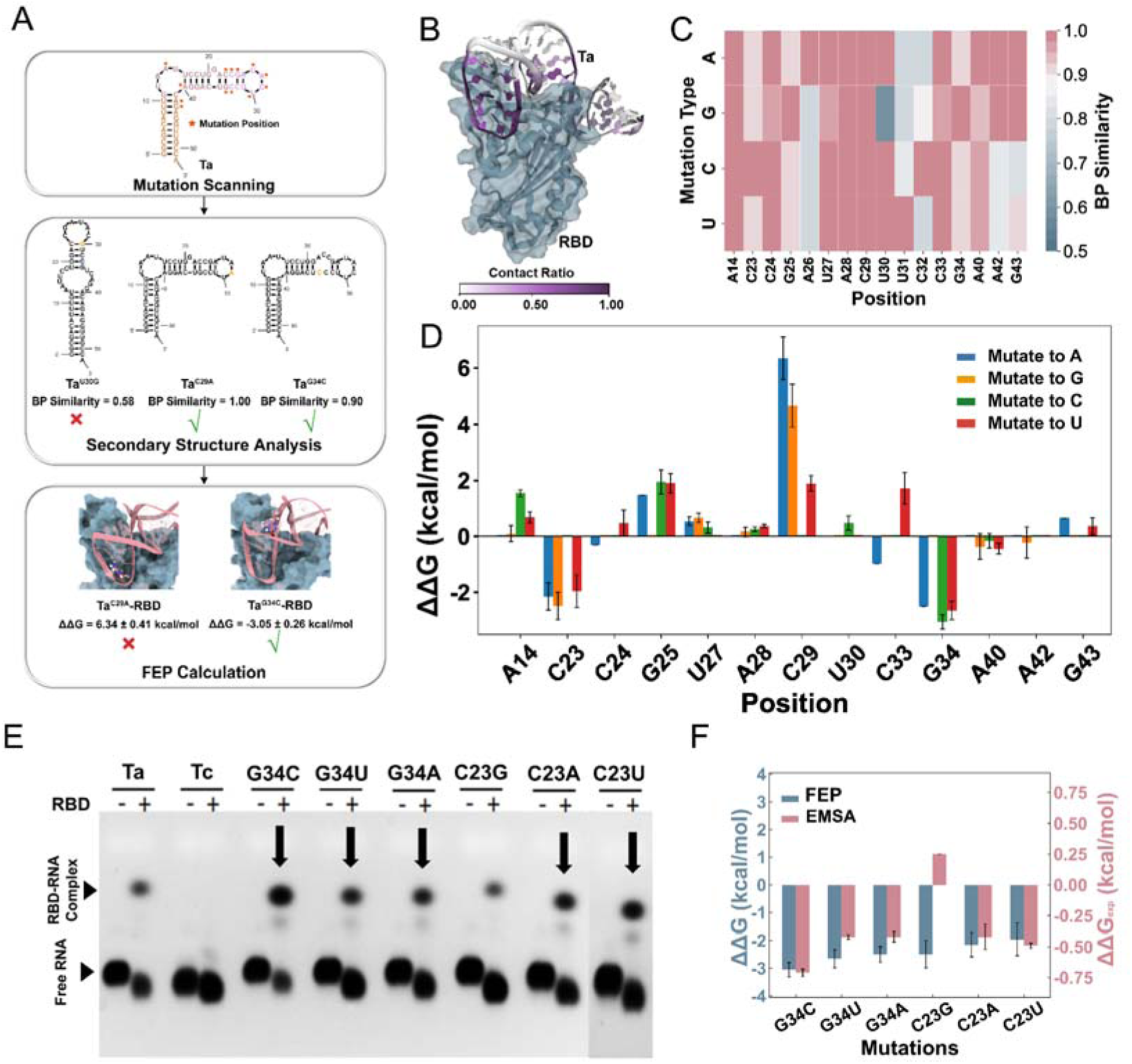
Structure-based rational design of Ta analogues with improved binding affinities and their experimental validation. (A) Flowchart illustrating the combination of rational mutation scanning, secondary structure analysis (SSA), and free energy perturbation (FEP) to optimize aptamer binding affinities. (B) Contact ratios of nucleotides on the aptamer Ta bound to RBD (dark blue surface). Data were collected from three independent MD simulations. (C) SSA based on BP similarity for 16 selected nucleotides mutated to other three bases. Definition of BP similarity was given in the main text. Only mutations with BP similarity greater than 0.9 were subjected to further FEP calculations. (D) The binding free energy changes assessed by FEP calculations for selected single mutations. Data represent mean±s.d. collected from five independent simulations. (E) EMSA results of Ta, Tc, and 6 designed candidate sequences bound to the RBD of SARS-CoV-2 spike protein. The aptamer-RBD complex bands were detected by running an agarose gel after incubation of 40 μM of RBD protein with 0.5 μM indicated aptamer variant. (F) Comparison of the binding free energy changes derived from FEP calculations (ΔΔ*G*, left scale) and EMSA experiments (ΔΔ*G_exp_*, right scale) for 6 designed candidate aptamers. Data represent mean±s.d. collected from 5 (FEP)/three (EMSA) independent replications.

Based on the binding complex between the aptamer Ta and RBD constructed above, we selected 16 vital nucleotides (A14, C23-G34, A40, and A42-G43) on the aptamer Ta that showed high contact frequency with RBD (see Fig. 4B) for exhaustive single-mutation scanning. These selected nucleotides play a crucial role in Ta-RBD binding interactions, and their mutagenesis studies may aid in designing analogues with improved binding affinity.

After SSA based on BP similarity, we found that most single mutations preserved a WT-like 2D structure (Fig. 4C) with BP similarity greater than 0.9 (such as Ta^G34C^ in Fig. S8), but several single mutations cause significant rearrangement of their 2D structure with a BP similarity less than 0.9 (for instance, Ta^U30G^ in Fig. S8). Mutations with a BP similarity greater than 0.9 were subjected to FEP calculations to further evaluate their impact on binding free energy changes, with the results shown in Fig. 4D and Table S3.

For nucleotides in the apical loop of the aptamer Ta (see Figs. 1B and 2E), mutation to any other base types resulted in an increased binding free energy and weakened binding affinity, such as ΔΔ*G*=1.94±0.43 kcal/mol for G25C, 0.66±0.16 kcal/mol for U27G, and 4.66±0.76 kcal/mol for C29G. This suggests that G25, U27 and C29 are already optimal choices for RBD binding. The previous study [16] highlighted the importance of the apical loop in directing the aptamer Ta’s binding to the RBD, and truncation of this apical loop was proven to significantly reduce its binding ability. Our FEP calculations align with this observation. Next, though the end stem part of the aptamer Ta also tightly binds to the RBD (see Figs. 2E and 4B), base mutations in this region had minimal effects on binding affinity, likely due to dominant electrostatic interactions between the negatively charged phosphate groups and the positively charged residues of RBD. For instance, A42G showed a ΔΔ*G of*-0.22±0.56 kcal/mol and G43U a ΔΔ*G* of 0.37±0.30 kcal/mol. Furthermore, since the bulge part of the aptamer Ta serves as a linker to regulate the saddle-like binding shape with RBD (see Figs. 2E and 4B), nucleotide mutations in this region may also affect binding ability despite their low contact frequency with the RBD. Indeed, disruption of the C23-G34 base pair improved the binding affinity of the aptamer to RBD, with C23A, C23G, and C23U yielding ΔΔ*G* values of-2.15±0.43,-2.49±0.49, and-1.96±0.58 kcal/mol, respectively, and G34A, G34C, and G34U showing ΔΔ*G* values of-2.50±0.28,-3.05± 0.26, and-2.65±0.33 kcal/mol, respectively (see Fig. 4D and Table S3). Ultimately, we generated six candidate sequences (Ta^C23A^, Ta^C23G^, Ta^C23U^, Ta^G34A^, Ta^G34C^, and Ta^G34U^) predicted to have stronger binding affinity to RBD than WT aptamer Ta.

We conducted EMSA experiments to validate the binding ability of these designed aptamers. Preliminary binding results of the RBD-RNA complex bands showed that five of the six designed sequences (Ta^G34C^, Ta^G34U^, Ta^G34A^, Ta^C23A^, and Ta^C23U^) exhibited higher fluorescence intensity than WT aptamer Ta (see Fig. 4E). Further, the binding curves with different RBD concentrations and the resulting dissociation constant (*K_d_*) measurements confirmed the superior binding capabilities of these five sequences compared to WT aptamer Ta (see Fig. S9 and Table S2). As shown in Fig. 4F, except for Ta^C23G^, the binding free energy changes derived from EMSA experiments (ΔΔ*G_exp_*) were consistent with FEP calculations (ΔΔ*G*) for the remaining five designed aptamers. Here, the experimental binding free energy change is ΔΔG_exp_ =-k_B_Tln(Kd^wt^⁄Kd^mut^), where Kd^wt^ and Kd^mut^ are dissociation constants for WT aptamer Ta and the designed sequence, respectively, and k_B_ is the Boltzmann constant and T = 310 K. The magnitude of the binding free energy changes generated from FEP calculations tend to be greater, likely due to limitations in the force field parameters. Nonetheless, the high success rate (0.83, 5/6) achieved in this structure-based rational design process underscores the reliability of the RBD-aptamer Ta complex model proposed in Fig. 2E. These results highlight the potential of our CAAMO framework as an effective tool for optimizing aptamer binding affinity.

### 2.5 Designed aptamer Ta^G34C^ shows excellent binding ability to RBD

Among the five designed candidate sequences, both the FEP calculation and EMSA experiment confirmed that aptamer Ta^G34C^ has the highest binding affinity to the RBD of SARS-CoV-2 spike protein. As shown in Fig. 5A, the dissociation constant derived from EMSA experiments for aptamer Ta^G34C^ is *K_d_*=33.5±1.6 µM, which is approximately 3.3-fold higher compared to WT Ta (*K_d_*=110.7 µM). The remaining four designed aptamers exhibited approximately a 2-fold increase in binding affinity relative to WT Ta, with *K_d_*=55.5±4.1 µM for Ta^G34A^ and *K_d_*=55.7±9.8 µM for Ta^C23A^ (see Fig. S9 and Table S2).

**Fig. 5.**
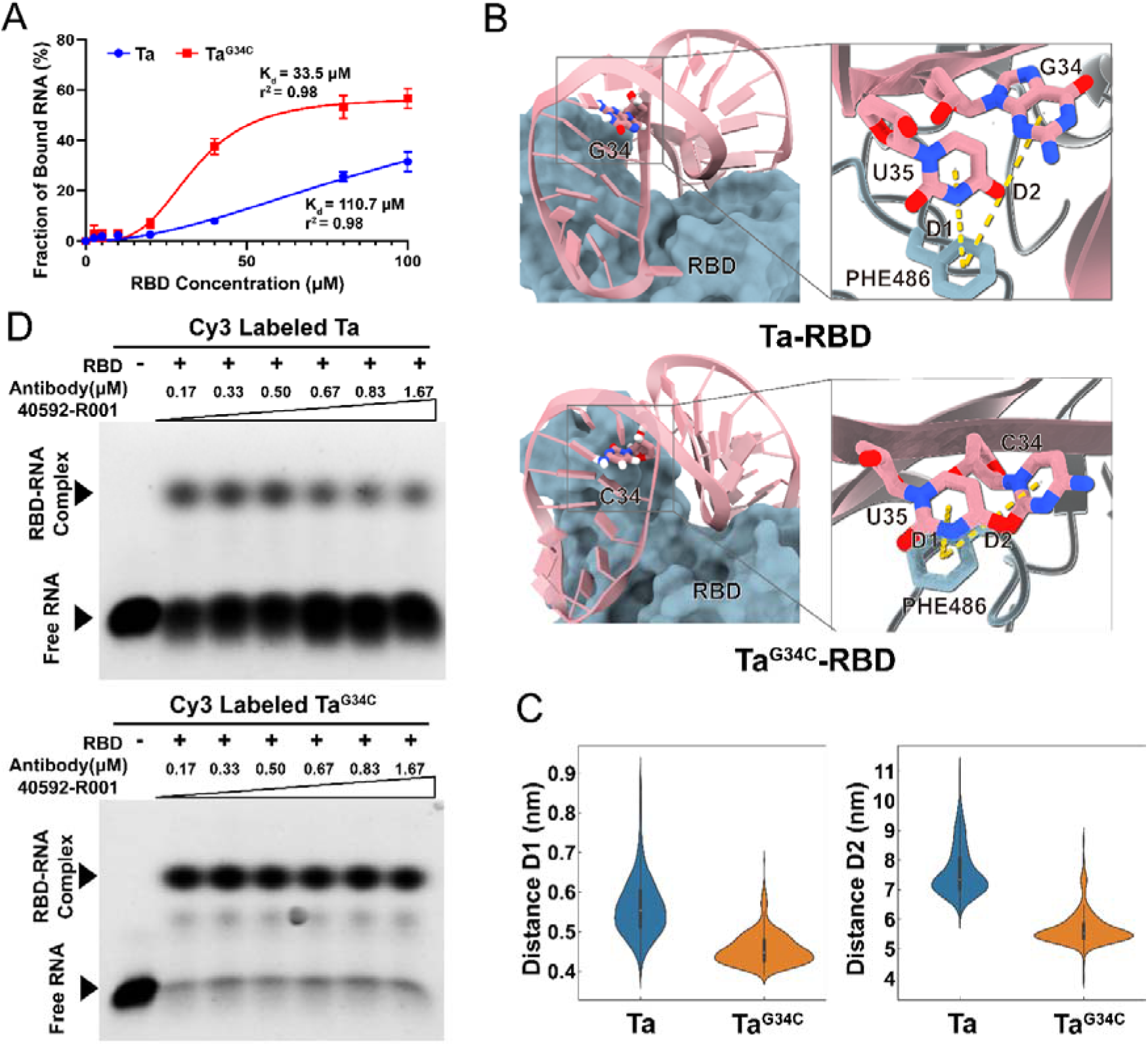
Designed aptamer Ta^G34C^ showed superior binding ability to WT Ta or antibody. (A) Binding curves and the resultant dissociation constants (*K_d_*) for WT Ta and the designed Ta^G34C^ with RBD protein.

We performed MD simulations on the Ta^G34C^-RBD binding complex to explore the molecular mechanism underlying the improved binding affinity of aptamer Ta^G34C^. As shown in the Fig. 5B, the C23-G34 base pair is located in the bulge region of WT Ta, which plays a critical role in regulating its saddle-like binding shape. We speculated that disrupting the C23-G34 base pair through mutation (e.g., G34C) could reduce the strain during aptamer binding to the RBD. The G34C substitution can alter the binding environment of adjacent nucleotides, such as U35, allowing them to form tighter contacts with the RBD loop region (from GLN474 to TYR489). For instance, our simulations showed that the distance between nucleotide U35 and PHE486 is shorter in the Ta^G34C^-RBD complex than that in the WT Ta-RBD complex (see Fig. 5C). This reduced distance remains stable throughout the MD simulations, indicating that U35 and PHE486 form a stable π-π stacking interaction after the G34C substitution. Additionally, the distance between C34 and PHE486 in Ta^G34C^-RBD is also closer compared to G34 and PHE486 in WT Ta-RBD. These findings support our hypothesis regarding the improved binding affinity in aptamer Ta^G34C^ and provide a basis for further in silico design of new aptamers.

We also conducted competitive binding experiments to compare the binding capacities of the designed aptamer Ta^G34C^ and a commercial monoclonal SARS-CoV-2 neutralizing antibody [39] against the RBD. As shown in Fig. 5D and Fig. 1F, when the monoclonal SARS-CoV-2 neutralizing antibody 40592-R001 was added to the WT Ta-RBD complex, the fluorescence intensity of the complex band gradually decreased, indicating that the antibody at high concentrations can partially replace the aptamer Ta in the Ta-RBD binding complex and has comparable, though weaker, binding ability. However, when the same antibody was added to the designed Ta^G34C^-RBD complex, the fluorescence intensity of the complex bands remained nearly unchanged at all tested antibody concentrations, indicating that the binding affinity of Ta^G34C^ is significantly stronger than that of the monoclonal SARS-CoV-2 neutralizing antibody or WT aptamer Ta.

To further exclude non-specific aptamer–protein interactions, we performed parallel EMSA assays using bovine serum albumin (BSA) as a non-target protein control for Ta, Tc, and the optimized Ta^G34C^ (see Fig. S10). Only weak, comparable background signals were observed for all three aptamers with BSA. Such minor non-specific binding may originate from BSA itself or trace contaminating proteins in the BSA samples (Fig. S10). In contrast, markedly stronger binding was detected between RBD and Ta or Ta^G34C^, whereas no detectable binding was observed with the negative control Tc (Figs. 4E, S10). Such distinct binding profiles of aptamers with RBD and BSA confirm that the aptamer–RBD interactions characterized in this study are target-specific.

In summary, the combined results from FEP calculated and EMSA measured binding affinities, binding molecular mechanism analysis, and antibody competitive binding assays clearly demonstrate that the designed aptamer Ta^G34C^ exhibits excellent binding ability to RBD. These findings highlight the importance of optimizing SELEX-derived aptamers through structure-based rational design to enhance their binding affinity.

The *K_d_* values were calculated from the EMSA image quantification with s.d. from three independent experiments (Mean±s.d., n = 3). (B) Comparison of the Ta-RBD and Ta^G34C^-RBD binding complexes. RNA is shown as a dark red ribbon while RBD is displayed as a dark blue surface. Zoom-in is the nucleotide G34 (WT Ta)/C34 (Ta^G34C^) and its surrounding nucleotide U35 and residue PHE486 shown in sticks. Atoms oxygen and nitrogen are colored in red and blue, respectively, and carbon atoms are colored by their locations. Definitions of two related distances (D1 and D2) are also displayed. (C) Violin plots of distributions of two related distances (D1 and D2) between selected RNA nucleotide-protein residue in Ta-RBD and Ta^G34C^-RBD binding complexes, respectively. Data were collected from three independent MD simulations. (D) EMSA images of competitive binding experiments to characterize the RBD binding abilities of RNA aptamers (WT Ta and Ta^G34C^) and the commercial monoclonal SARS-CoV-2 neutralizing antibody 40592-R001. The aptamer-RBD complex bands were showed by running an agarose gel after incubation of 40 μM of RBD protein and 0.5 μM indicated aptamer with varying concentrations of the antibody 40592-R001. Final antibody concentrations ranged from 0 to 1.67 μM in the reaction mixtures. Results showed that Ta^G34C^, but not WT Ta, exhibited a higher binding affinity to the RBD protein than that of the antibody.

### 2.6 Binding performance of Ta and Ta^G34C^ against SARS-CoV-2 RBD variants

To further evaluate the binding performance and specificity of the designed aptamer Ta^G34C^ toward different SARS-CoV-2 variants [40], we conducted extensive free energy perturbation combined with Hamiltonian replica-exchange molecular dynamics (FEP/HREX) [41–43] for both the wild-type aptamer Ta and the optimized Ta^G34C^ against a series of RBD mutants. The representative variants include the early Alpha (B.1.1.7) and Beta (B.1.351) lineages, as well as a panel of Omicron sublineages (BA.1–BA.5, BA.2.75, BQ.1, XBB, XBB.1.5, EG.5.1, HK.3, JN.1, and KP.3) carrying multiple mutations within the RBD region (residues 333–527). For each variant, mutations within 5 Å of the bound aptamer were included in the FEP to accurately estimate the relative binding free energy change (ΔΔ*G*).

For the wild-type Ta aptamer, the FEP-predicted binding affinities toward the Alpha and Beta RBD variants were consistent with the experimental trends, validating the reliability of our model (see Table S4). Specifically, Ta maintained comparable or slightly enhanced binding to the Alpha variant and showed only marginally reduced affinity for the Beta variant.

In contrast, the optimized aptamer Ta^G34C^ exhibited markedly improved and broad-spectrum binding toward most tested variants (see Table S5). For early variants such as Alpha, Beta, and Gamma, Ta^G34C^ maintained enhanced affinities (ΔΔ*G* < 0). Notably, for multiple Omicron sublineages—including BA.1, BA.2, BA.2.12.1, BA.2.75, XBB, XBB.1.5, XBB.1.16, XBB.1.9, XBB.2.3, EG.5.1, XBB.1.5.70, HK.3, BA.2.86, JN.1 and JN.1.11.1—the calculated binding free energy changes ranged from −1.89 to −7.58 kcal/mol relative to the wild-type RBD, indicating substantially stronger interactions despite the accumulation of multiple mutations at the aptamer–RBD interface. Only in a few other Omicron sublineages, such as BA.4, BA.5, and KP.3, a slight reduction in binding affinity was observed (ΔΔ*G* > 0).

These computational findings demonstrate that the Ta^G34C^ aptamer not only preserves high affinity and specificity for the RBD but also exhibits improved tolerance to the extensive mutational landscape of SARS-CoV-2. Collectively, our results suggest that Ta^G34C^ holds promise as a high-affinity and potentially cross-variant aptamer candidate for targeting diverse SARS-CoV-2 spike protein variants.

## 3. Discussion and conclusion

Determining the 3D structure of RNA aptamer-target protein complexes is crucial for understanding the binding mechanism and optimizing binding efficacy for various applications. Despite the discovery of aptamers for over 1,100 target proteins [44], only a limited number of aptamer-protein complex 3D structures are available (only 119 deposited in the PDB). The scarcity of experimentally determined aptamer-protein structures is primarily due to the inherent flexibility of RNA molecules and the high cost of experimental procedures. However, with recent advancement in protein and RNA 3D structure prediction [32,45,46] and improvements in atomic force field parameters [47,48], as well as the increased availability of high-performance computing resources, computational modeling of RNA aptamer-protein complexes has become increasingly promising. In this work, we developed a multi-strategy computational approach to determine the most probable RNA aptamer-protein binding conformation. Our approach integrates RNA 3D structure prediction, ensemble docking, clustering, and binding capacity assessment, with a critical focus on identifying the lowest-energy binding conformation using various energy assessment methods. We believe that the predicted binding conformation represents a plausible member of the predicted ensemble that is functionally informative for guiding structure-based aptamer optimization, although it may not correspond to the exact native structure.

We found that the aptamer Ta binds to the RBD of SARS-CoV-2 spike protein in a saddle-like shape, where the apical loop and end stem parts bind to two opposite sides of the RBD, while the bulge serves as a connector and regulates the bending angle. Theoretical analysis and experimental validation confirmed that this binding model is reliable. With this 3D structure, we were able to conduct structure-based rational optimization of the aptamer sequence to improve its binding affinity and then to perform a head-to-head comparison between the RNA aptamer and neutralizing antibodies.

The development of aptamer-based RNA therapeutics involves iterative rounds of design, testing, and optimization. A key step in this process is optimizing the original aptamer sequence to design a series of analogues with comparable or even improved binding affinities to the target protein. *In silico* structure-based aptamers design, which involves nucleotide mutation, insertion, deletion and their binding affinity assessment, is gaining prominence [49]. Our proposed framework, CAAMO, enables efficient aptamer lead optimization requiring only the aptamer’s nucleotide sequence as the input information. Unlike peptide and protein counterparts, RNA structures are highly sensitive to nucleotide composition, and even a single mutation may cause rearrangement in its secondary structure. To address this issue, we employed secondary structure analysis to evaluate the effect of mutations on RNA folding and only submitted those maintaining a similar folding structure to subsequent FEP-guided binding free energy evaluations.

For the Ta-RBD binding complex, mutation scanning revealed that mutations of nucleotides in direct contact with the RBD showed negligible or increased binding free energy changes, whereas mutations in the middle portion, which has fewer contacts with RBD but regulates the bending angle of RNA aptamer, resulted in reduced binding free energy and improved binding affinity. These findings suggest that during structure-based optimization, attention should be paid not only to the nucleotides at the binding interface but also to the regulatory nucleotides that affect the binding shape. Our CAAMO framework generated six optimized candidate sequences, and EMSA experiments confirmed that five of them exhibited significantly stronger binding affinity to the RBD than the wild-type Ta (about 2 to 3.3-fold improvement). This high success rate (∼83%) validates the reliability of the putative 3D binding model and demonstrates the effectiveness of our computational framework. Although the absolute *K_d_* values determined by EMSA cannot be directly compared with surface-based methods such as SPR or BLI, the relative affinity trends remain highly consistent. While EMSA provides semi-quantitative affinity estimates, the close agreement between experimental EMSA trends and FEP-calculated ΔΔ*G* values supports the robustness of the relative affinity changes reported here. In future studies, additional orthogonal biophysical techniques (e.g., filter-binding, SPR, or BLI) will be employed to further validate and refine the protein–aptamer interaction models. Furthermore, since the current study only considered single-nucleotide mutations, other design strategies such as double mutations, multiple mutations, nucleotide insertions and deletions should be explored in the future studies. Beyond the Ta–RBD system, the CAAMO framework itself is inherently generalizable. Our ongoing work applying CAAMO to optimize aptamers targeting other therapeutically relevant proteins, such as the epidermal growth factor receptor (EGFR) [50], has yielded promising preliminary results, further underscoring the potential of this computational–experimental integrated approach for broader aptamer engineering. While the present study primarily focused on affinity enhancement, we acknowledge that other key developability traits—such as nuclease resistance, structural and thermodynamic stability, and *in vivo* persistence—are equally critical for advancing aptamers toward therapeutic applications. These properties were not evaluated here but will be systematically addressed in future iterations of the CAAMO framework to enable comprehensive optimization of aptamer candidates.

Aptamers, often referred to as “chemical antibody”, offer a valuable comparison to real antibodies in terms of binding properties, yet such comparative analyses are currently limited. The RBD of SARS-CoV-2 spike protein is an ideal target for these comparisons, as binding modes for both the aptamer Ta and neutralizing antibody-RBD complexes are available. Our computational and experimental studies showed that the aptamer Ta has comparable binding abilities to the RBD compared to representative neutralizing antibodies analyzed in this study. Analysis of the underlying binding details revealed that more critical RBD residues are involved in aptamer Ta-RBD interaction, while MM/GBSA calculations and competitive binding assays confirmed the stronger binding capacity of the aptamer Ta. Specifically, a newly designed aptamer (Ta^G34C^) exhibited excellent RBD binding ability, surpassing both WT Ta and commercial antibodies. Given the concerns surrounding antibody-dependent enhancement (ADE), which could worsen COVID-19 outcomes by increasing virus infectivity and virulence [51], there is an urgent need to develop smaller and safer molecules to neutralize the virus and complement existing treatment modalities. Furthermore, the previous study [16] has claimed that WT Ta aptamer can efficiently block viral infection at low concentration and provide a promising lead for the detection and treatment of SARS-CoV-2 and emerging variants. Therefore, the designed aptamer Ta^G34C^, as well as other four successful candidates, provides a promising alternative to antibodies for fighting COVID-19.

## 4. Materials and Methods

**4.1 Generation of aptamer 3D conformation ensemble**

To predict the 3D structures of the aptamers Ta (5’-GGCGACAUUU GUAAUUCCUG GACCGAUACU UCCGUCAGGA CAGAGGUUGC CA-3’) and Tc (5’-GGUCCUGGAC CGAUACUUCC GUCAGGACCA-3’), five state-of-the-art RNA 3D structure prediction tools were employed: IsRNA2 [32], FARFAR2 [31], SimRNA [33], iFoldRNA [52], and RNAComposer [53]. The predicted secondary structure from Mfold [30], along with the aptamer sequence, were used as input for 3D structure prediction. IsRNA2 was downloaded and ran locally following the identical procedure as in the previous studies [54]; FARFAR2 was executed in Rosetta 3.12 using the *rna_denovo* application with default parameters; web-based version of SimRNA, iFoldRNA, and RNAComposer were assessed, and their default parameters were used. The top 5 predictions generated by each program (25 structures in total) were collected to construct a 3D conformation ensemble of a given RNA aptamer.

### 4.2 Generation of aptamer-RBD binding complex pool

The binding complex pool for aptamer Ta-RBD was generated using multiple popular docking tools, including HADDOCK [55], HDOCK [56], ZDOCK [57], and RosettaDock [58]. Rosettadock was performed locally using Rosetta software, while the other three docking tasks were completed on their respective webservers. The RBD structure [23] (PDB id: 6LZG) refined by MD simulation, and 25 RNA 3D conformations predicted in the previous step were used as the receptor and ligands for docking, respectively. The top 10 binding poses from each docking were recorded, yielding a total of 1,000 (4×25×10) aptamer Ta-RBD binding complexes in the pool. Of these, 879 conformations shared the same binding interface of ACE2-RBD complex. Then, these 879 conformations were clustered into 6 major groups, each representing a significantly distinct binding conformation. The binding complex pool for Tc-RBD interaction (used as a negative control) was constructed following the same procedure.

### 4.3 Molecular dynamics simulations

All-atom MD simulations were performed using Gromacs [59] (version 2021.5). For each system, a water box with at least 1.5 nm distance from the surface of the complex was used to solvate the systems, and NaCl ions were added to achieve a physiological concentration of 150 mM after neutralizing the system. The Amberff14SB force field was used for protein and RNA. Water molecules were described by TIP3P model [60] and Li and Merz ion parameters [61] were used. The periodic boundary conditions were applied in all three dimensions. The particle mesh Ewald (PME) method was used to compute the long-range electrostatic interactions while the vdW interactions were truncated at 1.5 nm. The LINCS algorithm was adopted to constrain the H-bonds to allow an integration timestep of 2 fs. Before MD productions, an energy minimization, 100 ps NVT, and 10 ns NPT simulations with a temperature of T = 310.15 K and pressure of 1 atm was executed sequentially to equilibrate the simulation box. To prevent unexpected structural deviations in the beginning, position restraints on backbone atoms of RNA and protein were performed in the NVT and NPT simulations. After that, series of MD simulations were conducted in the NPT ensemble with velocity-rescaled Berendsen thermostat: (1) 500ns MD simulations were performed for the RBD. Snapshots extracted from the last 300 ns MD trajectories (600 snapshots recorded in the duration of 500 ps) were clustered based on linkage method with a 0.2 nm RMSD cutoff to obtain the most probable receptor conformation for docking. (2) 100 ns simulations were performed on the representative Ta/Tc-RBD complex conformations obtained from the docking. 5 mM MgCl_2_ was added to the simulation system. These trajectories were used to calculate RMSD and MM/GBSA. (3) 100 ns simulations were performed on the ACE2-RBD complex (PDBID: 6LZG) and three antibody-RBD complexes (PDBID: 7CDI, 7DK4 and 7K4N). These also were used to perform MM/GBSA. (4) 500ns simulations were conducted for the Ta^G34C^-RBD and Ta-RBD complexes. 5 mM MgCl_2_ was added to the simulation system. A summary of MD simulations performed in this study was given in Table S1. The 3D structure models were rendered using UCSF ChimeraX (version 1.6.1) programs.

### 4.4 MM/GBSA calculations

In this study, binding energies (Δ*G*) of ACE2, aptamers (Ta and Tc), and several neutralizing antibodies to RBD were assessed using the end-point Molecular Mechanics Generalized Born Surface Area (MM/GBSA) method, implemented through the gmx_MMPBSA software [62]. Briefly, Δ*G* was calculated by summing up the changes in electro-static energies (Δ*E^ele^*), the van der Waals energies (Δ*E^vdW^*), the electrostatic solvation energy (Δ*G^GB^*, polar contribution), the nonpolar contribution (Δ*G^SA^*) between the solute and the continuum solvent, and conformational entropy (*–T*Δ*S*) upon ligand binding. The dielectric constants for the solute and solvent were set to 10 and 78.5, respectively. The OBC solvation model (igb = 8) was used. The interaction entropy method was used to calculate the conformational entropy (*–T*Δ*S*). Additionally, the lowest energy conformations from the complexes pool clustering were selected based on single-frame conformational MM/GBSA. The static complex structure was employed to calculate the enthalpy using MM/GBSA.

### 4.5 SMD and rupture works

Constant velocity steered molecular dynamics (SMD) was performed to calculate the rupture work required to separate the bound aptamer from the RBD. A steered velocity of 0.1 nm/ns was applied to the center-of-mass (COM) of the aptamer, while keeping the COM of RBD fixed, using the COM distance between aptamer (or antibody) and RBD as the collective variable. Four replicate SMD simulations (about 50 ns simulation time each) were performed for each binding complex. All simulation parameters remain the same as those used in the MD simulations. The force spectra were recorded during the SMD simulations at a 0.1 ps intervals. The rupture works were obtained through integration of the force spectra over the COM distance.

### 4.6 FEP calculations

The binding free energy changes resulting from point mutations of key bases at the interface between the aptamer Ta and RBD were calculated using the free energy perturbation (FEP) method. We estimated the binding free energy changes for single nucleotide mutation in both the bound state (Ta and RBD complex) Δ*G_bound_* and the free state (the aptamer Ta only) Δ*G_free_* using Gromacs 2021.5. Thus, the binding free energy change caused by nucleotide mutation was estimated as ΔΔ*G_calc_*=Δ*G_bound_*-Δ*G_free_*. For each single nucleotide mutation, the dual-topology file was prepared in a pmx-like manner based on the Amberff14SB force field and eighteen λ windows (0.0, 0.01, 0.05, 0.1, 0.15, 0.2, 0.25, 0.35, 0.45, 0.55, 0.65, 0.75, 0.8, 0.85, 0.9, 0.95, 0.99, and 1.0) with 1 ns/window were used. The vdW and electrostatic interactions were transferred simultaneously during simulations and the soft-core potentials (α=0.3) were used. For each mutation, five independent replicas starting from different conditions were performed for sufficient sampling and at least 180 ns (1 ns×18 windows×5 runs×2 states) simulation time was generated, which result in reasonable convergence in the free energy calculations. The Gromacs bar analysis tool was used to estimate the binding free energy changes.

### 4.7 FEP/HREX

To evaluate the binding sensitivity of the optimized aptamer Ta^G34C^ toward SARS-CoV-2 RBD variants, we employed free energy perturbation combined with Hamiltonian replica-exchange molecular dynamics (FEP/HREX) simulations for enhanced sampling efficiency and improved convergence. The relative binding free energy changes (ΔΔ*G*) upon RBD mutations were estimated as:

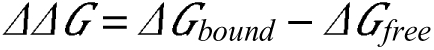

where Δ*G_bound_*and Δ*G_free_*represent the RBD mutations-induced free energy changes in the complexed and unbound states, respectively. All simulations were performed using GROMACS 2021.5 with the Amber ff14SB force field. For each mutation, dual-topology structures were generated in a pmx-like manner, and 32 λ-windows (0.0, 0.01, 0.02, 0.03, 0.06, 0.09, 0.12, 0.16, 0.20, 0.24, 0.28, 0.32, 0.36, 0.40, 0.44, 0.48, 0.52, 0.56, 0.60, 0.64, 0.68, 0.72, 0.76, 0.80, 0.84, 0.88, 0.91, 0.94, 0.97, 0.98, 0.99, 1.0) were distributed uniformly between 0.0 and 1.0. To ensure sufficient sampling, each window was simulated for 5 ns, with five independent replicas initiated from distinct velocity seeds. Replica exchange between adjacent λ states was attempted every 1 ps to enhance phase-space overlap and sampling convergence. The van der Waals and electrostatic transformations were performed simultaneously, employing a soft-core potential (α = 0.3) to avoid singularities. For each RBD variant system, this setup resulted in an accumulated simulation time of approximately 1600 ns (5 ns × 32 windows × 5 replicas × 2 states). The Gromacs bar analysis tool was used to estimate the binding free energy changes.

### 4.8 EMSA experiments

The sequences of Ta, Tc, Ta^G34C^, Ta^G34U^, Ta^G34A^, Ta^C23G^, Ta^C23A^, and Ta^C23U^ aptamers are shown in Table S2. In these sequences, the uppercase letters A, U, G and C indicated the ribonucleotide. The EMSA was performed according to a previous protocol [54] with a minor modification. Synthesized 5′ end Cy3-labelled RNAs were resuspended with the RNase-free H_2_O to a concentration of 100 µM. 5 μl of Cy3-labelled RNAs were annealed with the 5 μl 2×annealing buffer (20 mM Tris-HCl pH 7.5, 200 mM KCl) under a predefined procedure: 68°C for 5min, then annealing at-0.1°C/s to 25°C, and finally at 25°C for 5 min, then diluted to the final concentration of 5 µM. The SARS-CoV-2 Spike RBD protein was purchased from HuaBio (Cat: HA210064). Mouse monoclonal SARS-CoV-2 neutralizing antibody was purchased from Sino Biological (Cat: 40592-R001). 40 μM of RBD protein and 0.5 μM of annealed Cy3-labelled aptamers were mixed in the EMSA buffer (10 mM Sodium phosphate buffer pH 7.5, 1 U/μl SUPERase-In RNase Inhibitor [Thermo Fisher]). For competitive binding experiments, Cy3-labelled RNAs, RBD protein, and neutralizing antibody 40592-R001 were added simultaneously to the EMSA buffer and incubated at room temperature for 20 min.

The integrity and purity of the RBD protein were confirmed by denaturing SDS-PAGE (Fig. S11), showing a single intact band without degradation. The multiple bands observed in native PAGE (e.g., Fig. 3E) are due to conformational and glycosylation heterogeneity [63–66] rather than protein degradation. To rule out non-specific aptamer–protein interactions, BSA was additionally included as a non-target protein control in EMSA assays; the wild-type Ta, the negative control Tc, and the optimized Ta^G34C^ all showed only weak, comparable background signals with BSA but distinct target-specific binding to RBD (Fig. S10). Uncropped EMSA gel images (Fig. S12) and consistent results from three biological replicates (Fig. 2F and S9) confirm the absence of protein aggregation and ensure data reliability.

For competitive binding experiments, 40 μM of RBD protein, 0.5 μM of annealed Cy3-labelled RNAs and increasing concentrations of SARS-CoV-2 neutralizing antibody 40592-R001 (0–1.67 μM) were mixed in the EMSA buffer and incubated at room temperature for 20 min. Next, the mixtures were added 6×loading buffer (15% Ficoll 400, 0.25% Bromophenol Blue, 0.25% Xylene cyanol, 1×TBE), then resolved on a native 0.8% agarose gel and imaged with iBright1500 (Thermo Fisher). The images were quantified with Image J software. The dissociation constant *K_d_* for RBD with aptamers were calculated using Prism 8 (GraphPad) software. All uncropped raw gel images corresponding to these EMSA experiments are provided in Supplementary Fig. S12.

## Acknowledgments

We thank Zaixing Yang and Teng Xie for their helpful discussions. This work was supported by funds from the National Key R&D Program of China (2021YFF1200404 to R.Z., 2021YFA1201200 to R.Z.), National Natural Science Foundation of China (Nos. 12474203 to D.Z., U1967217 to R.Z.), National Independent Innovation Demonstration Zone Shanghai Zhangjiang Major Projects (ZJZX2020014 to R.Z.), the Zhejiang Provincial Natural Science Foundation of China (No. LZ25A040001), Starry Night Science Fund at Shanghai Institute for Advanced Study of Zhejiang University (SN-ZJU-SIAS-003 to R.Z., SN-ZJU-SIAS-006 to L.H.), National Center of Technology Innovation for Biopharmaceuticals (NCTIB2022HS02010 to R.Z.), Shanghai Artificial Intelligence Lab (P22KN00272 to R.Z.), Aoming Biomedical Research (AO-ZJU-SIAS-001 to R.Z.), and Zhejiang University Global Partnership Fund (188170+194452409/004 to R.Z.).

## CRediT authorship contribution statement

R.Z. and D.Z. designed research; Y.Y. and Y.J. performed all molecular simulations; L.Q. performed EMSA experiments under the supervision of Z.W.; Y.Y. contributed new reagents/analytic tools; Y.Y., Z.W., D.Z., and R.Z. analyzed data; Y.Y., L.Q., Z.W., D.Z., D.B., L.H., and R.Z. wrote the paper with contributions from all authors.

