## Supplemental Tables and Figures for "*In silico* design and validation of high-affinity RNA aptamers for SARS-CoV-2 comparable to neutralizing antibodies"

**Table.** S1. Summary of simulation systems in this study.

| **Name** | **Number of**  **atoms** | **Production simulation time** | **Description** |
| --- | --- | --- | --- |
| Ta (3D) | 1106 | 50 ns×10 replicas replica exchange MD, 2 runs | Predict RNA 3D structure in the coarse-grained IsRNA2 model for Ta aptamer |
| Tc (3D) | 636 | 50 ns×10 replicas replica exchange MD, 2 runs | Predict RNA 3D structure in the coarse-grained IsRNA2 model for Tc aptamer |
| RBD | 62442 | 500ns, 3 runs | Relax the 3D structure of RBD in all-atom MD simulations to prepare docking structure |
| RBD&Ta  (Conf 01) | 130136 | 500ns, 3 runs | Relax the 3D binding complex of RBD & Ta from initial conformation |
| RBD&Ta  (Conf 02) | 130136 | 500ns, 3 runs | Relax the 3D binding complex of RBD & Ta from initial conformation |
| RBD&Ta  (Conf 03) | 285645 | 500ns, 3 runs | Relax the 3D binding complex of RBD & Ta from initial conformation |
| RBD&Ta  (Conf 04) | 217783 | 500ns, 3 runs | Relax the 3D binding complex of RBD & Ta from initial conformation |
| RBD&Ta  (Conf 05) | 179307 | 500ns, 3 runs | Relax the 3D binding complex of RBD & Ta from initial conformation |
| RBD&Ta  (Conf 06) | 120009 | 500ns, 3 runs | Relax the 3D binding complex of RBD & Ta from initial conformation |
| RBD&Ta  (Conf 01)  (MM/GBSA) | 130136 | 100ns, 3 runs | Simulate the binding complex of Ta and RBD to perform MM/GBSA |
| RBD&Ta  (Conf 02)  (MM/GBSA) | 130136 | 100ns, 3 runs | Simulate the binding complex of Ta and RBD to perform MM/GBSA |
| RBD&Ta  (Conf 03)  (MM/GBSA) | 285645 | 100ns, 3 runs | Simulate the binding complex of Ta and RBD to perform MM/GBSA |
| RBD&Ta  (Conf 04)  (MM/GBSA) | 217783 | 100ns, 3 runs | Simulate the binding complex of Ta and RBD to perform MM/GBSA |
| RBD&Ta  (Conf 05)  (MM/GBSA) | 179307 | 100ns, 3 runs | Simulate the binding complex of Ta and RBD to perform MM/GBSA |
| RBD&Ta  (Conf 06)  (MM/GBSA) | 120009 | 100ns, 3 runs | Simulate the binding complex of Ta and RBD to perform MM/GBSA |
| RBD&Ta  (Conf 01)  (SMD) | 284617 | ~50ns, 4 runs | Separate the bound Ta from the RBD to perform SMD |
| RBD&Ta  (Conf 02)  (SMD) | 130136 | ~50ns, 4 runs | Separate the bound Ta from the RBD to perform SMD |
| RBD&Ta  (Conf 03)  (SMD) | 285645 | ~50ns, 4 runs | Separate the bound Ta from the RBD to perform SMD |
| RBD&Ta  (Conf 04)  (SMD) | 217783 | ~50ns, 4 runs | Separate the bound Ta from the RBD to perform SMD |
| RBD&Tc  (Conf 01) | 60549 | 500ns, 3 runs | Relax the 3D binding complex of RBD & Tc from initial conformation |
| RBD&Tc  (Conf 02) | 51210 | 500ns, 3 runs | Relax the 3D binding complex of RBD & Tc from initial conformation |
| RBD&Tc  (Conf 03) | 47448 | 500ns, 3 runs | Relax the 3D binding complex of RBD & Tc from initial conformation |
| RBD&Tc  (Conf 04) | 50101 | 500ns, 3 runs | Relax the 3D binding complex of RBD & Tc from initial conformation |
| RBD&Tc  (Conf 01)  (MM/GBSA) | 60549 | 100ns, 3 runs | Simulate the binding complex of Tc and RBD to perform MM/GBSA |
| RBD&Tc  (Conf 02)  (MM/GBSA) | 51210 | 100ns, 3 runs | Simulate the binding complex of Tc and RBD to perform MM/GBSA |
| RBD&Tc  (Conf 03)  (MM/GBSA) | 47448 | 100ns, 3 runs | Simulate the binding complex of Tc and RBD to perform MM/GBSA |
| RBD&Tc  (Conf 04)  (MM/GBSA) | 50101 | 100ns, 3 runs | Simulate the binding complex of Tc and RBD to perform MM/GBSA |
| RBD&Tc  (Conf 01)  (SMD) | 60549 | ~50ns, 4 runs | Separate the bound Tc from the RBD to perform SMD |
| RBD&Tc  (Conf 02)  (SMD) | 51210 | ~50ns, 4 runs | Separate the bound Tc from the RBD to perform SMD |
| RBD&Tc  (Conf 03)  (SMD) | 47448 | ~50ns, 4 runs | Separate the bound Tc from the RBD to perform SMD |
| RBD&Tc  (Conf 04)  (SMD) | 50101 | ~50ns, 4 runs | Separate the bound Tc from the RBD to perform SMD |
| RBD&ACE2 | 117622 | 500ns, 3 runs | Simulate the 3D binding complex of RBD & ACE2 to calculate contact ratios |
| RBD&P2C-1F11 | 254496 | 500ns, 3 runs | Relax the 3D binding complex of RBD & P2C-1F11 from initial conformation |
| RBD&2H2 Fab | 364742 | 500ns, 3 runs | Relax the 3D binding complex of RBD & 2H2 Fab from initial conformation |
| RBD&S2E12 Fab | 291645 | 500ns, 3 runs | Relax the 3D binding complex of RBD & S2E12 Fab from initial conformation |
| RBD&ACE2  (MM/GBSA) | 117622 | 100ns, 3 runs | Simulate the binding complex of ACE2 and RBD to perform MM/GBSA |
| RBD&P2C-1F11  (MM/GBSA) | 254496 | 100ns, 3 runs | Simulate the binding complex of P2C-1F11 and RBD to perform MM/GBSA |
| RBD&2H2 Fab  (MM/GBSA) | 364742 | 100ns, 3 runs | Simulate the binding complex of 2H2 Fab and RBD to perform MM/GBSA |
| RBD&S2E12 Fab  (MM/GBSA) | 291645 | 100ns, 3 runs | Simulate the binding complex of S2E12 Fab and RBD to perform MM/GBSA |
| Ta (free) | 129115 | 500ns, 3 runs | Relax the 3D structure of Ta in all-atom MD simulations to prepare FEP free state |
| RBD&Ta^G34C^ | 108541 | 500ns, 3 runs | Simulate the binding complex of Ta^G34C^ mutation & RBD |

**Table.** S2. The aptamer sequences employed in this study and their binding energies with RBD.

| **Aptamer Name** | **Sequences** | ***K_d_* (µM)** | ***ΔΔG_exp_* (kcal/mol)** | ***ΔΔG_FEP_***  **(kcal/mol)** |
| --- | --- | --- | --- | --- |
| Ta | 5’-GGCGACAUUU  GUAAUUCCUG  GACCGAUACU  UCCGUCAGGA  CAGAGGUUGCCA-3’ | 110.7 | -- | -- |
| Tc | 5’-GGUCCUGGAC  CGAUACUUCC  GUCAGGACCA-3’ | -- | -- | -- |
| Ta^G34C^ | 5’- GGCGACAUUU  GUAAUUCCUG  GACCGAUACU  UCCCUCAGGA  CAGAGGUUGCCA-3’ | 33.5±1.6 | -0.71±0.03 | -3.05±0.26 |
| Ta^G34U^ | 5’- GGCGACAUUU  GUAAUUCCUG  GACCGAUACU  UCCUUCAGGA  CAGAGGUUGCCA-3’ | 54.7±2.0 | -0.42±0.02 | -2.65±0.33 |
| Ta^G34A^ | 5’- GGCGACAUUU  GUAAUUCCUG  GACCGAUACU  UCCAUCAGGA  CAGAGGUUGCCA-3’ | 55.5±4.1 | -0.42±0.04 | -2.50±0.08 |
| Ta^C23G^ | 5’- GGCGACAUUU  GUAAUUCCUG  GAGCGAUACU  UCCGUCAGGA  CAGAGGUUGCCA-3’ | 167.9 | 0.25±0.00 | -2.49±0.49 |
| Ta^C23A^ | 5’-GGCGACAUUU  GUAAUUCCUG  GAACGAUACU  UCCGUCAGGA  CAGAGGUUGCCA-3’ | 55.7±9.8 | -0.42±0.10 | -2.15±0.43 |
| Ta^C23U^ | 5’- GGCGACAUUU  GUAAUUCCUG  GAUCGAUACU  UCCGUCAGGA  CAGAGGUUGCCA-3’ | 48.7±2.0 | -0.49±0.02 | -1.96±0.58 |

**Table.** S3. The relative binding free energy changes for single nucleotide mutations on Ta binding. Mean ± standard deviation (kcal/mol) from five independent FEP runs are given.

| **Mutation** | ***ΔG^bond^*** | ***ΔG^free^*** | ***ΔΔG^calc^*** |
| --- | --- | --- | --- |
| A14G | -53.12±0.27 | -53.22±0.06 | 0.10±0.28 |
| A14C | -81.05±0.11 | -82.58±0.07 | 1.54±0.13 |
| A14U | 18.99±0.19 | 18.31±0.05 | 0.68±0.20 |
| C23A | 84.01±0.20 | 86.16±0.38 | -2.15±0.43 |
| C23G | 30.35±0.24 | 32.84±0.42 | -2.49±0.49 |
| C23U | 102.20±0.45 | 104.16±0.37 | -1.96±0.58 |
| C24A | 82.30±0.18 | 82.61±0.16 | -0.30±0.24 |
| C24U | 99.57±0.43 | 99.11±0.18 | 0.47±0.47 |
| G25A | 57.94±0.37 | 56.48±0.34 | 1.47±0.50 |
| G25C | -22.95±0.41 | -24.89±0.14 | 1.94±0.43 |
| G25U | 78.16±0.24 | 76.26±0.25 | 1.90±0.35 |
| U27A | -18.65±0.04 | -19.19±0.09 | 0.54±0.10 |
| U27G | -70.85±0.09 | -71.51±0.13 | 0.66±0.16 |
| U27C | -99.46±0.18 | -99.78±0.08 | 0.32±0.20 |
| A28G | -52.81±0.13 | -52.97±0.12 | 0.16±0.17 |
| A28C | -82.67±0.06 | -82.92±0.05 | 0.26±0.08 |
| A28U | 17.93±0.04 | 17.55±0.05 | 0.37±0.06 |
| C29A | 89.42±0.30 | 83.08±0.28 | 6.34±0.41 |
| C29G | 34.47±0.75 | 29.91±0.13 | 4.66±0.76 |
| C29U | 100.81±0.11 | 98.91±0.16 | 1.89±0.28 |
| U30A | -20.35±0.12 | -20.35±0.05 | 0.00±0.13 |
| U30C | -99.91±0.19 | -100.39±0.18 | 0.48±0.26 |
| C33A | 81.44±0.46 | 82.41±0.17 | -0.97±0.49 |
| C33U | 99.98±0.48 | 98.26±0.30 | 1.72±0.56 |
| G34A | 55.91±0.15 | 58.41±0.24 | -2.50±0.28 |
| G34C | -27.51±0.14 | -24.46±0.21 | -3.05±0.26 |
| G34U | 74.17±0.24 | 76.82±0.22 | -2.65±0.33 |
| A40G | -55.18±0.41 | -54.82±0.21 | -0.36±0.46 |
| A40C | -81.47±0.17 | -81.31±0.20 | -0.16±0.26 |
| A40U | 19.69±0.08 | 20.12±0.17 | -0.44±0.19 |
| A42G | -53.07±0.49 | -52.85±0.26 | -0.22±0.56 |
| G43A | 52.94±0.26 | 52.30±0.15 | 0.65±0.30 |
| G43U | 75.93±0.11 | 75.57±0.27 | 0.37±0.30 |

**Table.** S4. Relative binding free energy changes (*ΔΔG*, kcal/mol) for Ta binding to SARS-CoV-2 RBD variants (Alpha and Beta) calculated by FEP/HREX. Experimental values (*ΔΔG_exp_*) are taken from Liu et al., PNAS 2021, doi: 10.1073/pnas.2112942118. FEP-calculated results (*ΔΔG_FEP_*) represent the mean ± standard deviation from five independent simulations.

| **Variants** | **Mutations** | ***ΔΔG_exp_*** | ***ΔΔG_FEP_*** |
| --- | --- | --- | --- |
| 20I (Alpha, V1) (B.1.1.7) | **N501Y** | -0.24 | -0.42±0.07 |
| 20H (Beta, V2) (B.1.351) | **K417N, E484K, N501Y** | 0.36 | 0.64±0.25 |

**Table.** S5. Relative binding free energy changes (*ΔΔG*, kcal/mol) for Ta^G34C^ binding to SARS-CoV-2 RBD variants calculated by FEP/HREX. The “Mutations” column lists all amino acid mutations within the RBD region (residues 333–527). Mutations located within 5 Å of the aptamer are highlighted in red and were explicitly perturbed in the FEP calculations. Values represent the mean ± standard deviation from five independent runs.

| **Variants** | **Mutations** | ***ΔΔG_FEP_*** |
| --- | --- | --- |
| 20I (Alpha, V1) (B.1.1.7) | **N501Y** | -0.81±0.08 |
| 20H (Beta, V2) (B.1.351) | **K417N, E484K, N501Y** | -0.67±0.26 |
| 20J (Gamma, V3) (P.1) | **K417T, E484K, N501Y** | -0.72±0.25 |
| 21K (Omicron) (BA.1) | **G339D, S371L, S373P, S375F, K417N, N440K, G446S, S477N, T478K, E484A, Q493R, G496S, Q498R, N501Y, Y505H** | -3.00±0.52 |
| 21L (Omicron) (BA.2) | **G339D, S371F, S373P, S375F, T376A, D405N, R408S, K417N, N440K, S477N, T478K, E484A, Q493R, Q498R, N501Y, Y505H** | -2.54±0.60 |
| 22A (Omicron) (BA.4) | **G339D, S371F, S373P, S375F, T376A, D405N, R408S, K417N, N440K, L452R, S477N, T478K, E484A, F486V, Q498R, N501Y, Y505H** | 2.11±0.67 |
| 22B (Omicron) (BA.5) | **G339D, S371F, S373P, S375F, T376A, D405N, R408S, K417N, N440K, L452R, S477N, T478K, E484A, F486V, Q498R, N501Y, Y505H** | 2.27±0.68 |
| 22C (Omicron) (BA.2.12.1) | **G339D, S371F, S373P, S375F, T376A, D405N, R408S, K417N, N440K, L452Q, S477N, T478K, E484A, Q493R, Q498R, N501Y, Y505H** | -2.40±0.41 |
| 22D (Omicron) (BA.2.75) | **G339H, S371F, S373P, S375F, T376A, D405N, R408S, K417N, N440K, G446S, N460K, S477N, T478K, E484A, Q493R, Q498R, N501Y, Y505H** | -5.03±0.81 |
| 22E (Omicron) (BQ.1) | **G339D, S371F, S373P, S375F, T376A, D405N, R408S, K417N, N440K, K444T, L452R, N460K, S477N, T478K, E484A, F486V, Q493R, Q498R, N501Y, Y505H** | -0.26±0.75 |
| 22F (Omicron) (XBB) | **G339H, R346T, L368I, S371F, S373P, S375F, T376A, D405N, R408S, K417N, N440K, V445P, G446S, N460K, S477N, T478K, E484A, F486S, F490S, Q493R, Q498R, N501Y, Y505H** | -3.13±0.73 |
| 23A (Omicron) (XBB.1.5) | **G339H, R346T, L368I, S371F, S373P, S375F, T376A, D405N, R408S, K417N, N440K, V445P, G446S, N460K, S477N, T478K, E484A, F486P, F490S, Q493R, Q498R, N501Y, Y505H** | -2.28±0.96 |
| 23B (Omicron) (XBB.1.16) | **G339H, R346T, L368I, S371F, S373P, S375F, T376A, D405N, R408S, K417N, N440K, V445P, G446S, N460K, S477N, T478R, E484A, F486P, F490S, Q493R, Q498R, N501Y, Y505H** | -2.64±0.79 |
| 23C (Omicron) (CH.1.1) | **G339H, R346T, S371F, S373P, S375F, T376A, D405N, R408S, K417N, N440K, K444T, G446S, L452R, N460K, S477N, T478K, E484A, F486S, Q493R, Q498R, N501Y, Y505H** | -0.80±0.95 |
| 23D (Omicron) (XBB.1.9) | **G339H, R346T, L368I, S371F, S373P, S375F, T376A, D405N, R408S, K417N, N440K, V445P, G446S, N460K, S477N, T478K, E484A, F486S, F490S, Q493R, Q498R, N501Y, Y505H** | -4.34±0.68 |
| 23E (Omicron) (XBB.2.3) | **G339H, R346T, L368I, S371F, S373P, S375F, T376A, D405N, R408S, K417N, N440K, V445P, G446S, N460K, S477N, T478K, E484A, F486P, F490S, Q493R, Q498R, N501Y, Y505H** | -3.28±1.06 |
| 23F (Omicron) (EG.5.1) | **G339H, R346T, L368I, S371F, S373P, S375F, T376A, D405N, R408S, K417N, N440K, V445P, G446S, F456L, N460K, S477N, T478K, E484A, F486P, F490S, Q493R, Q498R, N501Y, Y505H** | -5.32±1.38 |
| 23G (Omicron) (XBB.1.5.70) | **G339H, R346T, L368I, S371F, S373P, S375F, T376A, D405N, R408S, K417N, N440K, V445P, G446S, L455F, F456L, N460K, S477N, T478K, E484A, F486P, F490S, Q493R, Q498R, N501Y, Y505H** | -4.23±0.77 |
| 23H (Omicron) (HK.3) | **G339H, R346T, L368I, S371F, S373P, S375F, T376A, D405N, R408S, K417N, N440K, V445P, G446S, L455F, F456L, N460K, S477N, T478K, E484A, F486P, F490S, Q493R, Q498R, N501Y, Y505H** | -4.63±0.96 |
| 23I (Omicron) (BA.2.86) | **G339H, K356T, S371F, S373P, S375F, T376A, R403K, D405N, R408S, K417N, N440K, V445H, G446S, N450D, L452W, N460K, S477N, T478K, N481K, E484K, F486P, Q493R, Q498R, N501Y, Y505H** | -1.89±1.27 |
| 24A (Omicron) (JN.1) | **G339H, K356T, S371F, S373P, S375F, T376A, R403K, D405N, R408S, K417N, N440K, V445H, G446S, N450D, L452W, L455S, N460K, S477N, T478K, N481K, E484K, F486P, Q493R, Q498R, N501Y, Y505H** | -7.58±0.86 |
| 24B (Omicron) (JN.1.11.1) | **G339H, K356T, S371F, S373P, S375F, T376A, R403K, D405N, R408S, K417N, N440K, V445H, G446S, N450D, L452W, L455S, F456L, N460K, S477N, T478K, N481K, E484K, F486P, Q493R, Q498R, N501Y, Y505H** | -4.66±1.45 |
| 24C (Omicron) (KP.3) | **G339H, K356T, S371F, S373P, S375F, T376A, R403K, D405N, R408S, K417N, N440K, V445H, G446S, N450D, L452W, L455S, F456L, N460K, S477N, T478K, N481K, E484K, F486P, Q493E, Q498R, N501Y, Y505H** | 3.48±1.14 |

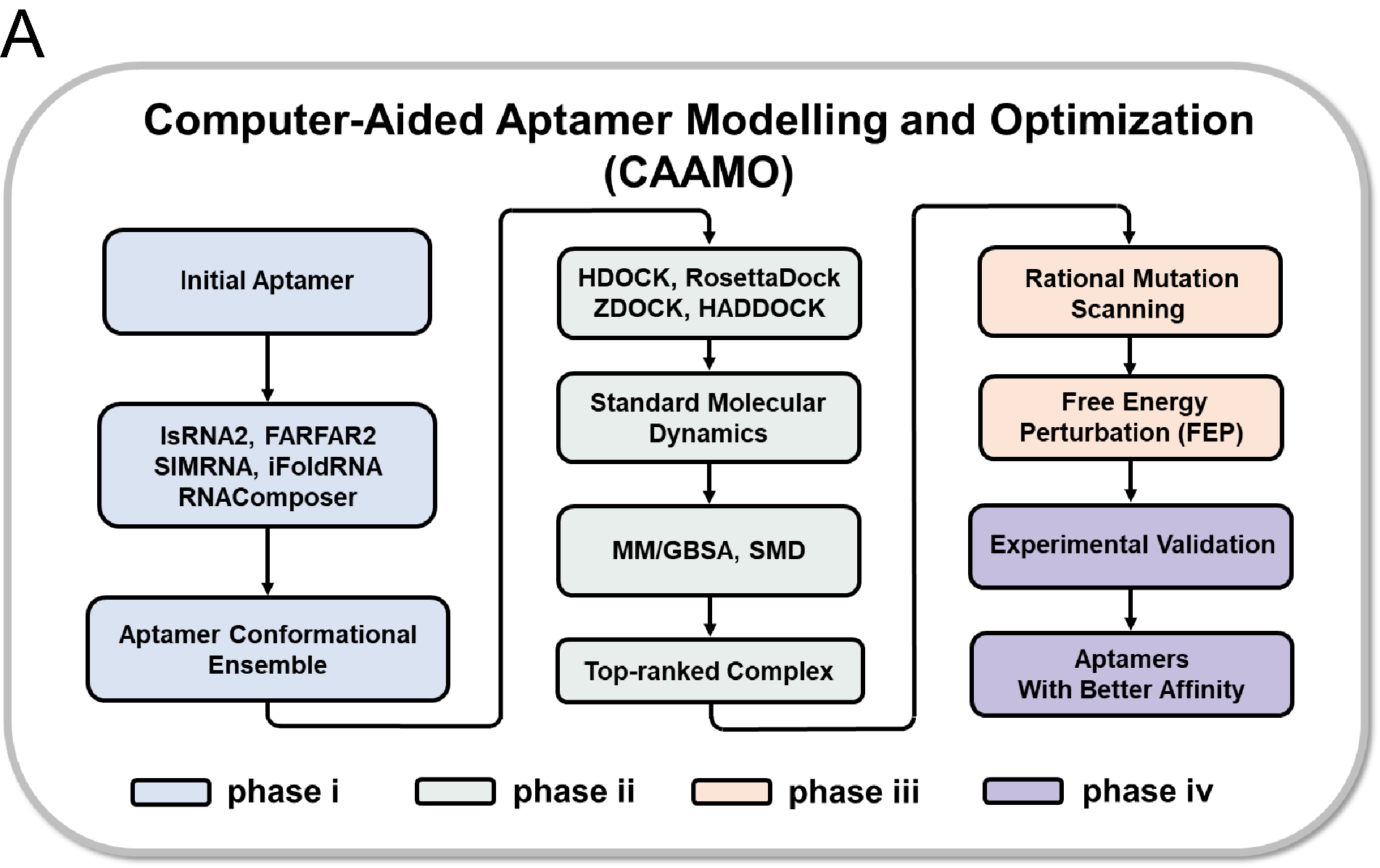

**Fig.** S1. The workflow of CAAMO. The workflow is to design high-affinity aptamers with physics-based simulations. It consists of four phases, including (i) constructing an aptamer conformational ensemble, (ii) identifying a proper aptamer binding mode, (iii) FEP-based rational design, and (iv) experimental validation.

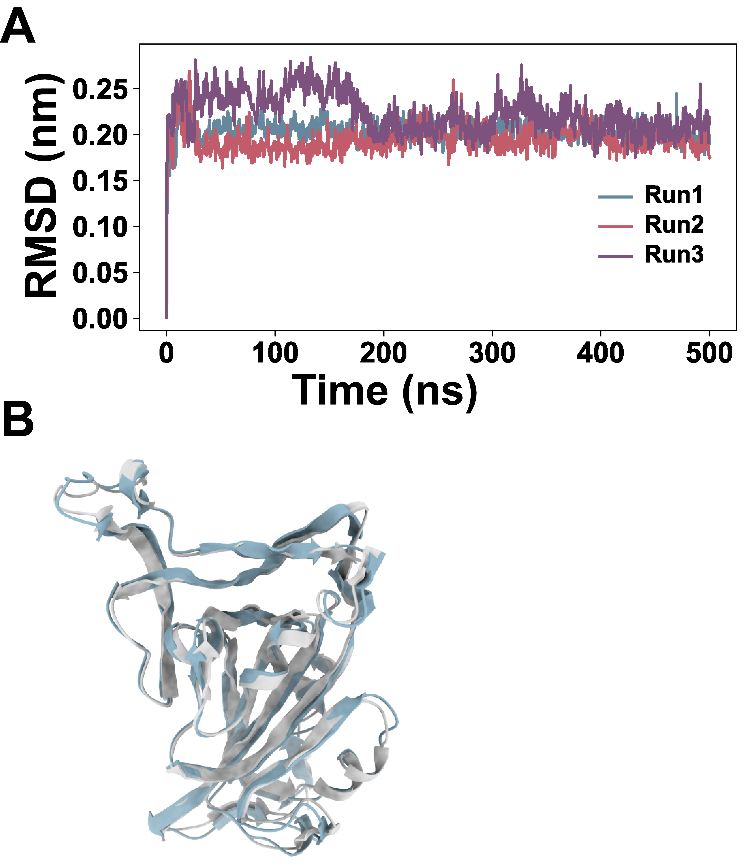

**Fig.** S2. (A) Heavy-atom RMSDs as functions of simulation time for RBD protein. Three 500-ns independent runs (Run1, Run2, and Run3) are performed. (B) Superposition of the MD refined structure (blue) and crystallographically resolved structure (PDB id: 6LZG, gray).

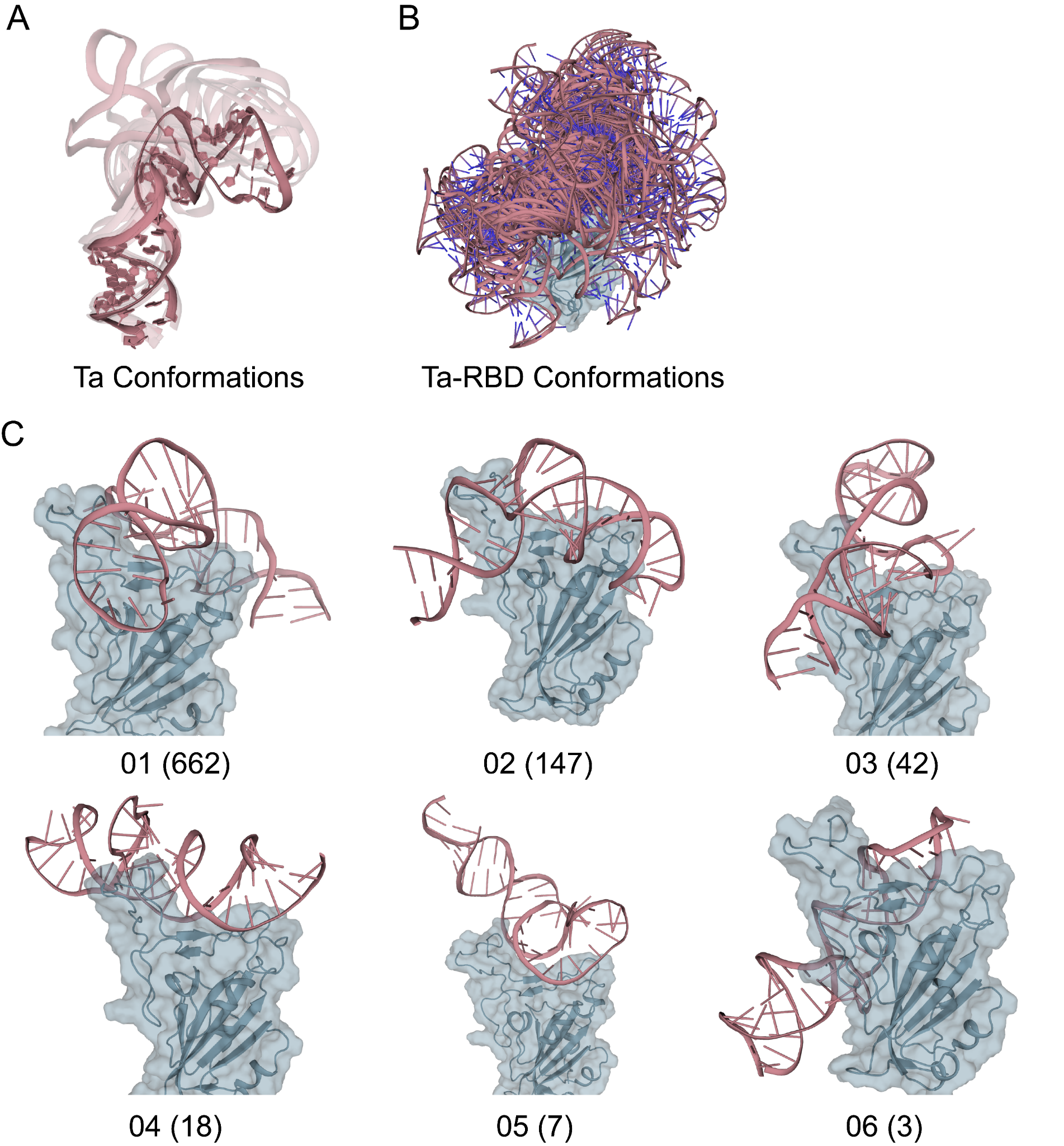

**Fig.** S3. 3D conformations of Ta and Ta-RBD complexes. (A) 25 Ta 3D structures were predicted by five RNA 3D modeling tools (IsRNA, FARFAR2, SimRNA, iFoldRNA and RNAcomposer). (B) The Ta-RBD complexes were predicted by four docking tools (HADDOCK, HDOCK, ZDOCK and RosettaDock). (C) Representative complex conformations from the top-six clusters are shown. The number of conformations contained within each cluster is indicated in parentheses.

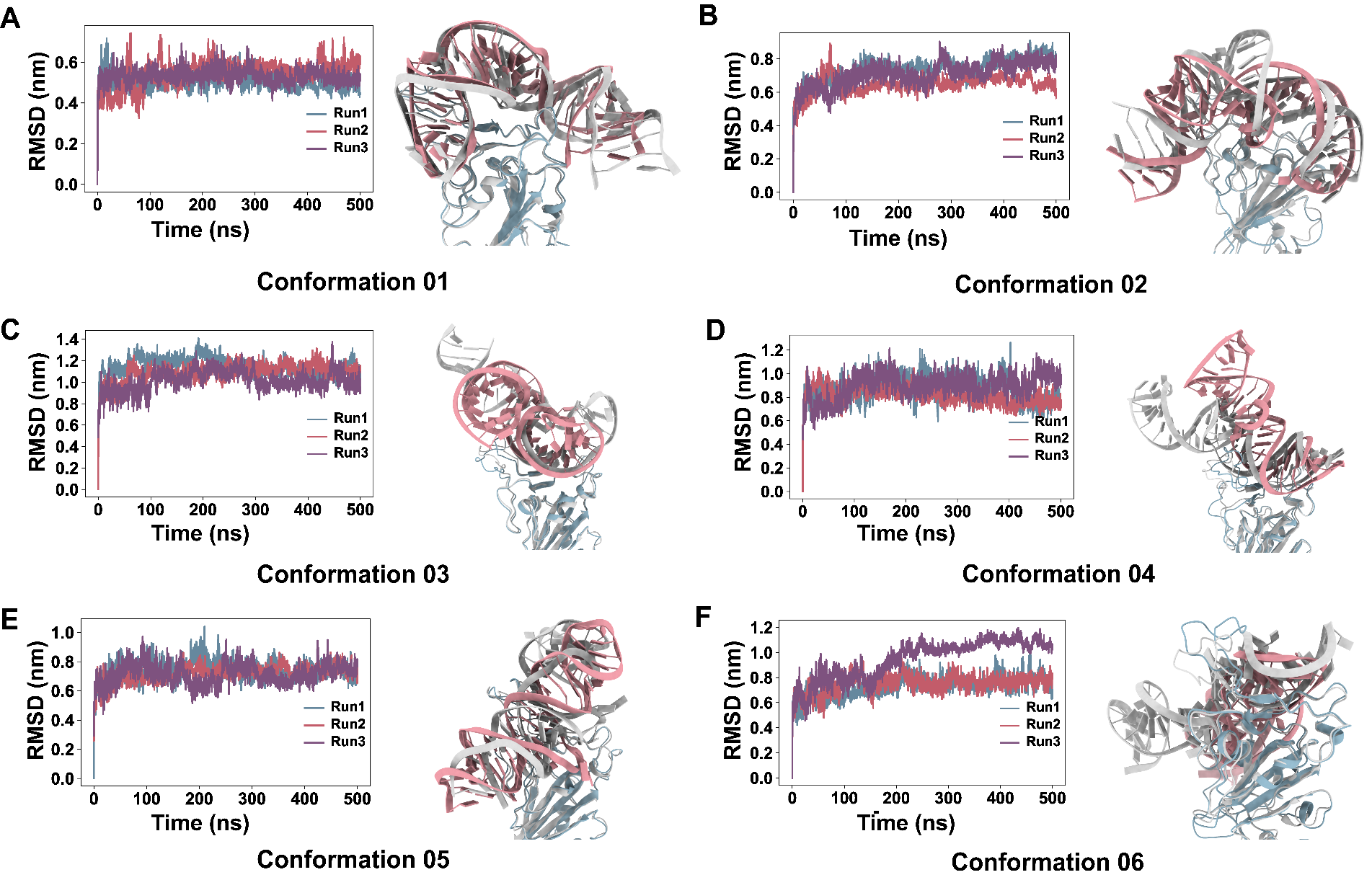

**Fig.** S4. RMSDs of heavy atoms are plotted as functions of simulation time across six selected conformations (Conformation 01-06) of Ta aptamer and RBD protein. Three independent 500-ns runs (Run1, Run2, and Run3) are performed. The initial structures are shown in white, while the MD refined structures are shown in red and blue for the aptamer and protein, respectively.

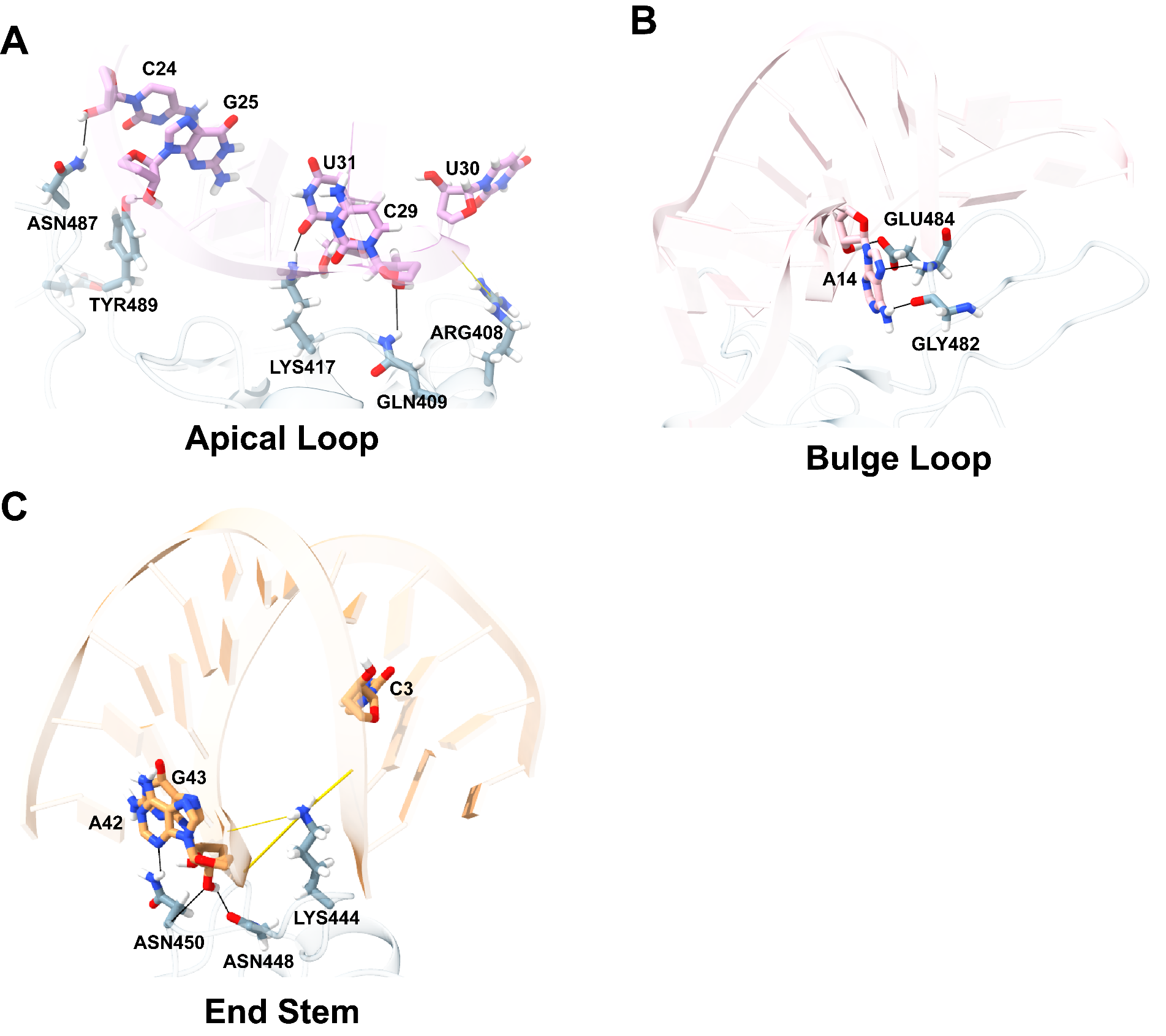

**Fig.** S5. The binding interfaces between the Ta aptamer and RBD. (A) Details of interactions between RBD and nucleotides C24, G25, C29, U30, and U31 in the apical loop part are presented. (B) Details of interactions between RBD and nucleotides A14 in the bulge loop. (C) Details of interactions between RBD and nucleotides C3, A42, and G43 in the end stem part. Hydrogen bonds between RNA nucleotides and RBD residues are indicated by thin black lines. Electrostatic interactions between RNA nucleotides and RBD residues are indicated by thin yellow lines.

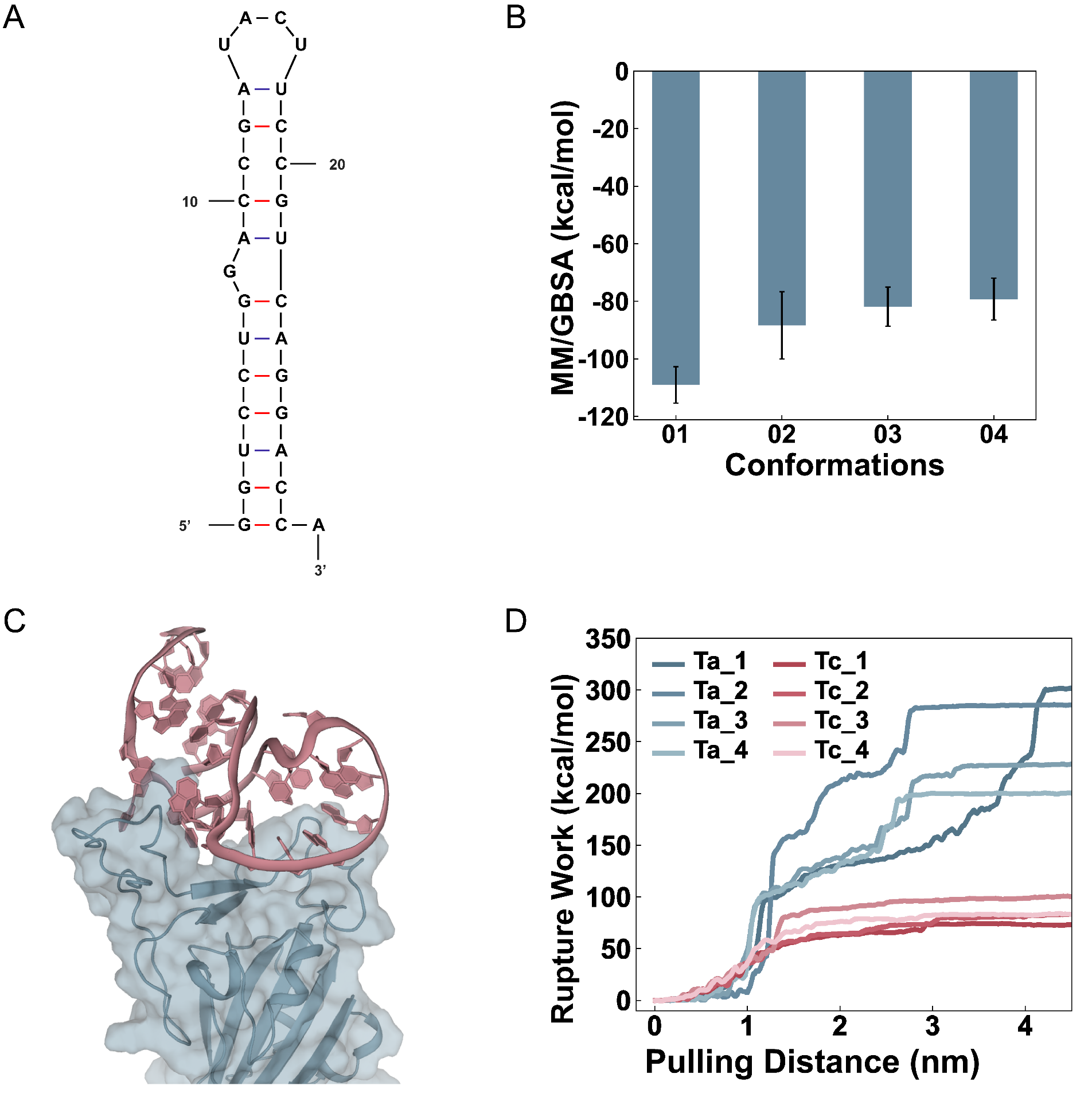

**Fig.** S6. The modeling of the negative control Tc-RBD complex. (A) The Mfold-predicted secondary structure of Tc. (B) The binding energies (*ΔG*) of four representative Tc-RBD conformations calculated by the MM/GBSA method. (C) The putative binding mode of Tc aptamer and RBD complex (D) The works required to pull the aptamer lead Ta and the negative control Tc away from RBD respectively. Ta-1 through Ta-4 and Tc-1 through Tc-4 represent four parallel replicates for the Ta and Tc simulations, respectively.

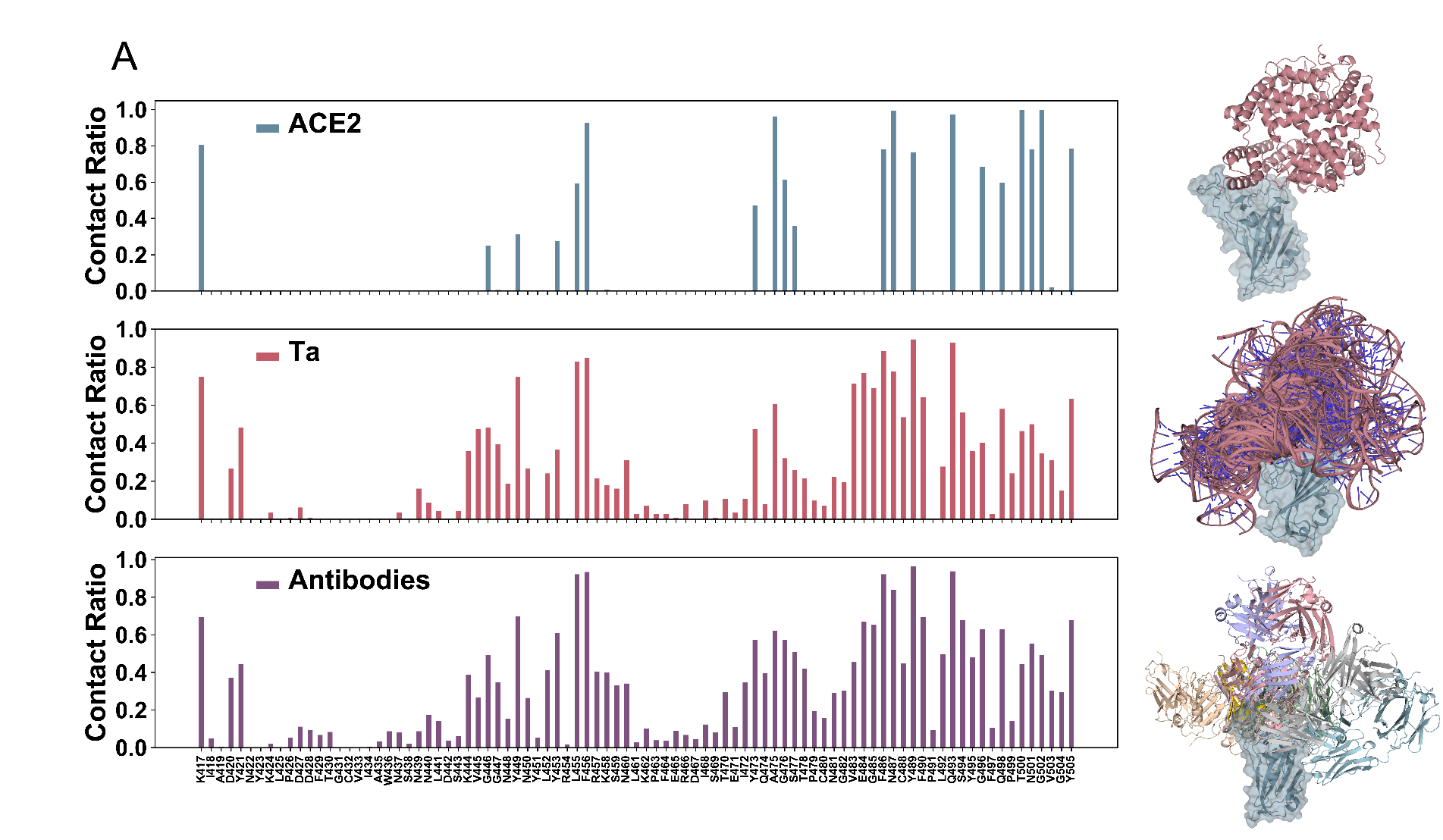

**Fig.** S7. Comparative analysis of RBD binding from ACE2, Ta and antibodies. MD simulations are performed on the ACE2-RBD complex and the contact ratios of RBD residues are obtained (upper panel). Subsequently, the contact ratios are determined for RBD residues in the Ta-RBD complexes (middle panel). Additionally, the contact ratios are accessed for RBD residues among all antibodies that bind to the ACE2-RBD interface, as recorded in the RCSB database (lower panel).

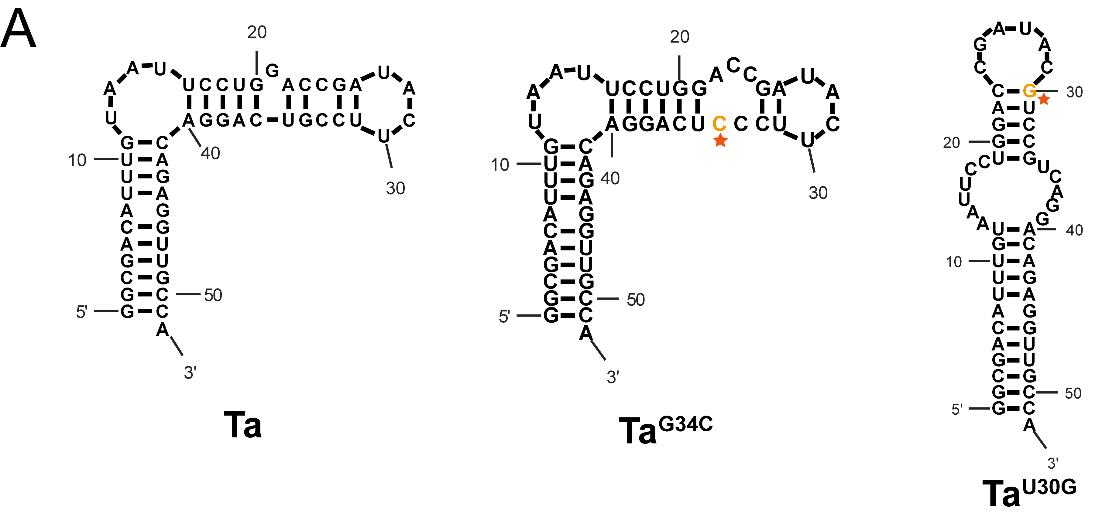

**Fig.** S8. The secondary structures of Ta, Ta^G34C^, and Ta^U30G^, predicted by Mfold. The mutated bases are highlighted in orange and marked with red pentagrams.

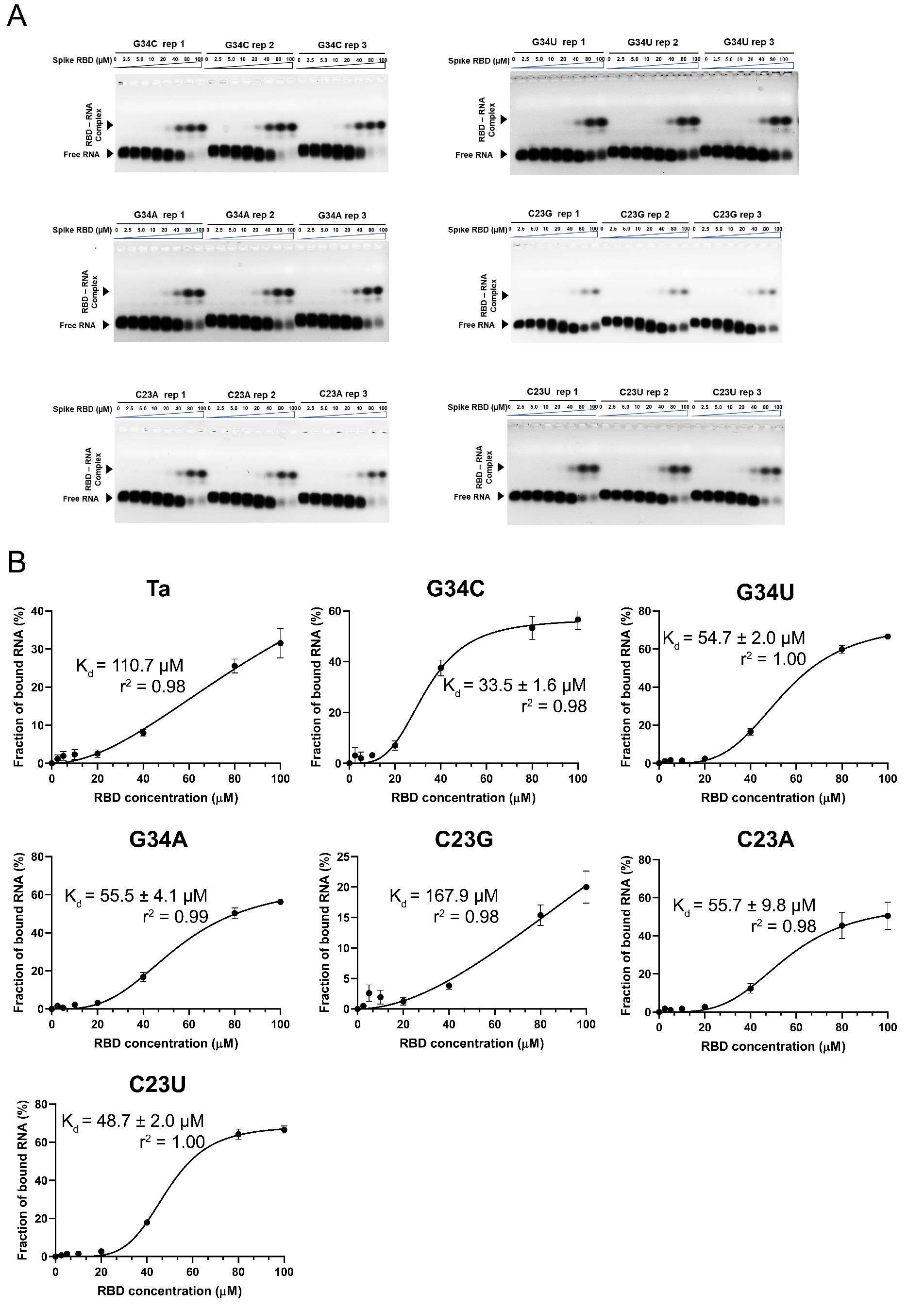

**Fig.** S9. (A) EMSA results of six Ta mutations (Ta^G34C^, Ta^G34U^, Ta^G34A^, Ta^C23G^, Ta^C23A^, and Ta^C23U^). (B) Binding curves and K_d_ values for Ta, Ta^G34C^, Ta^G34U^, Ta^G34A^, Ta^C23G^, Ta^C23A^, and Ta^C23U^. The K_d_ was calculated from the EMSA image quantification from three independent experiments (Mean ± s.d., n = 3).

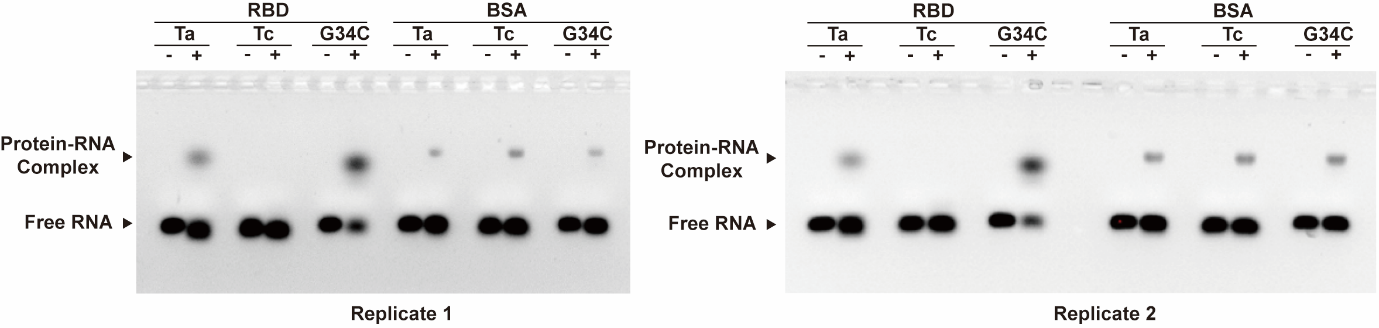

**Fig.** S10. EMSA assays with BSA as a non-target protein control verify the target-specific binding of designed aptamers to RBD. EMSA images of Ta, Tc, and Ta^G34C^ incubated with SARS-CoV-2 spike RBD or BSA protein. Aptamer–protein complex bands were visualized by agarose gel electrophoresis after incubation of 0.5 μM of the indicated aptamer with 40 μM RBD or BSA. Only weak, comparable background signals were observed with BSA for all three aptamers, whereas markedly stronger binding was detected between RBD and Ta or Ta^G34C^, while no detectable binding was observed between Tc and RBD, confirming that the aptamer–RBD interactions are target-specific.

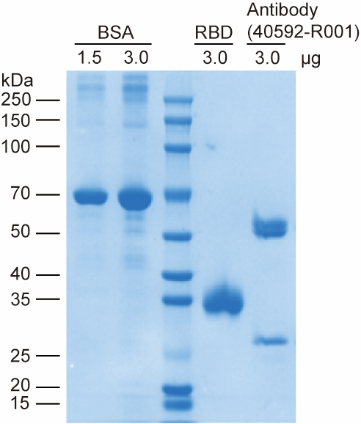

**Fig.** S11. SDS-PAGE analysis of the SARS-CoV-2 Spike RBD protein, neutralizing antibody (40592-R001) and BSA reference. This gel validates the high purity and structural integrity of the commercially sourced RBD protein and neutralizing antibody used in this study.

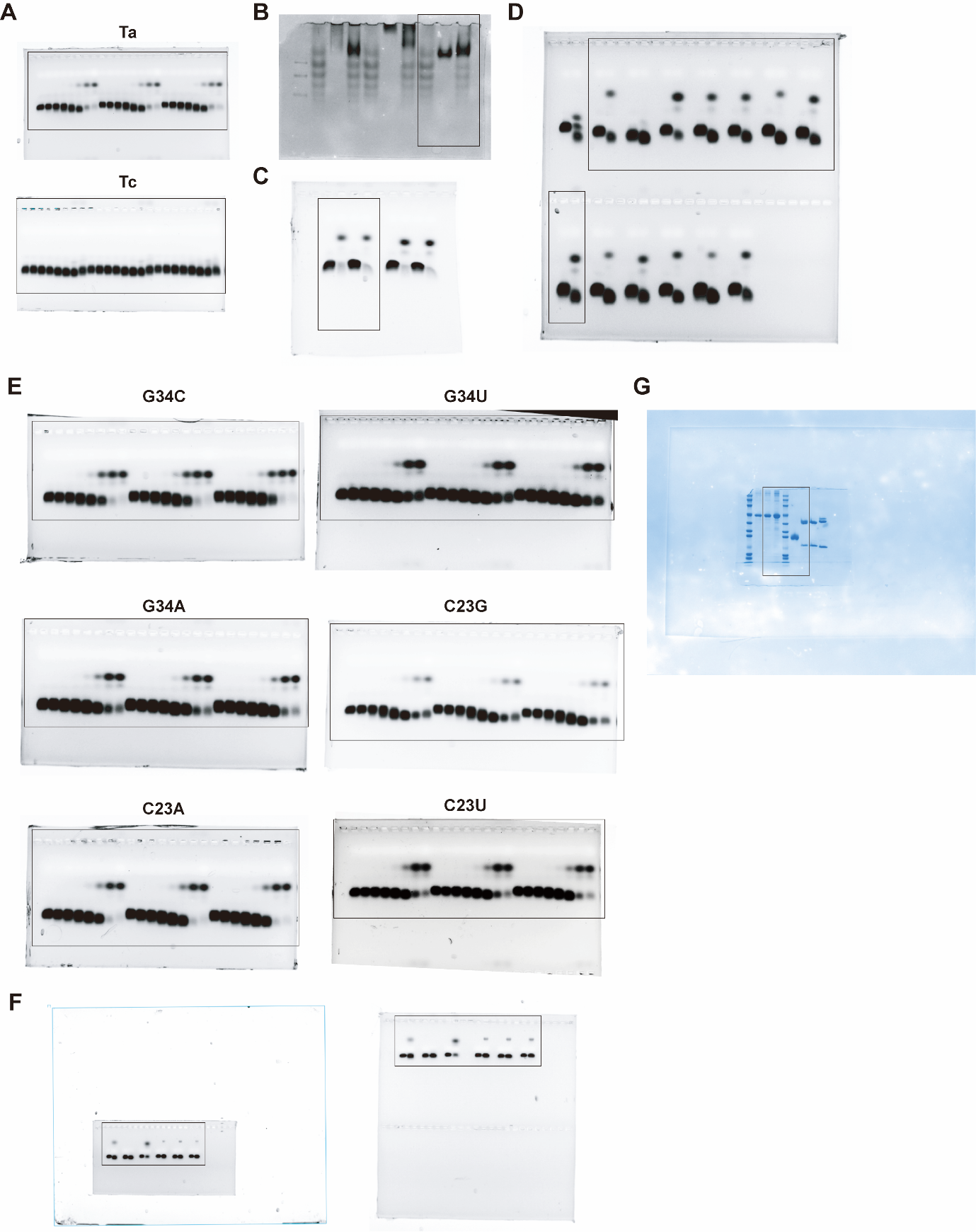

**Fig.** S12. Uncropped electrophoresis images corresponding to Figures 2F (A), 3D (B), 3E (C), 4E (D), S9A (E), S10 (F) and S11 (G). Black boxes indicate the cropped regions.
